# A hydrophobic core stabilizes the residual structure in the RRM2 intermediate state of the ALS-linked protein TDP-43

**DOI:** 10.1101/2024.06.12.598648

**Authors:** Brian C. Mackness, Brittany R. Morgan, Laura M. Deveau, Sagar V. Kathuria, Jill A. Zitzewitz, Francesca Massi

## Abstract

Folding intermediates mediate both protein folding and the misfolding and aggregation observed in human diseases, including amyotrophic lateral sclerosis (ALS), and are prime targets for therapeutic interventions. In this study, we identified the core nucleus of structure for a folding intermediate in the second RNA recognition motif (RRM2) of the ALS-linked RNA-binding protein, TDP-43, using a combination of experimental and computational approaches. Urea equilibrium unfolding studies revealed that the RRM2 intermediate state consists of collapsed residual secondary structure localized to the N-terminal half of RRM2, while the C-terminus is largely disordered. Steered molecular dynamics simulations and mutagenesis studies yielded key stabilizing hydrophobic contacts that, when mutated to alanine, severely disrupt the overall fold of RRM2. In combination, these findings suggest a role for this RRM intermediate in normal TDP-43 function as well as serving as a template for misfolding and aggregation through the low stability and non-native secondary structure.

## Introduction

Mutations in TDP-43 have been genetically linked to amyotrophic lateral sclerosis (ALS), providing direct evidence for the role of TDP-43 in disease pathogenesis (1). TDP-43 has also been associated with a host of other neurodegenerative diseases (1, 2), suggesting a common mechanism for TDP-43-mediated toxicity. A majority of patients with these TDP-43 proteinopathies lack disease-causing mutations, suggesting that misfolded or dysfunctional conformations of the wild-type protein are sufficient for the initiation and propagation of the disease phenotypes (3). Despite the plethora of evidence from cellular and animal models, the molecular basis of disease is not yet fully understood (4, 5).

TDP-43 is an RNA-binding protein containing two RNA recognition motifs (RRM1 and RRM2, Fig. 1A). It is primarily localized in the nucleus and is involved in numerous RNA processes (6). Evidence suggests that both loss of functional nuclear TDP-43 and accumulation of cytoplasmic aggregates are implicated in disease pathogenesis (4, 5). The disordered C-terminus and RRM2 are critical domains for the propagation of TDP-43 aggregation and toxicity in cell culture models (7), suggesting these domains may adopt pathogenic conformations that lead to cellular toxicity and neurodegeneration.

**Figure 1.**
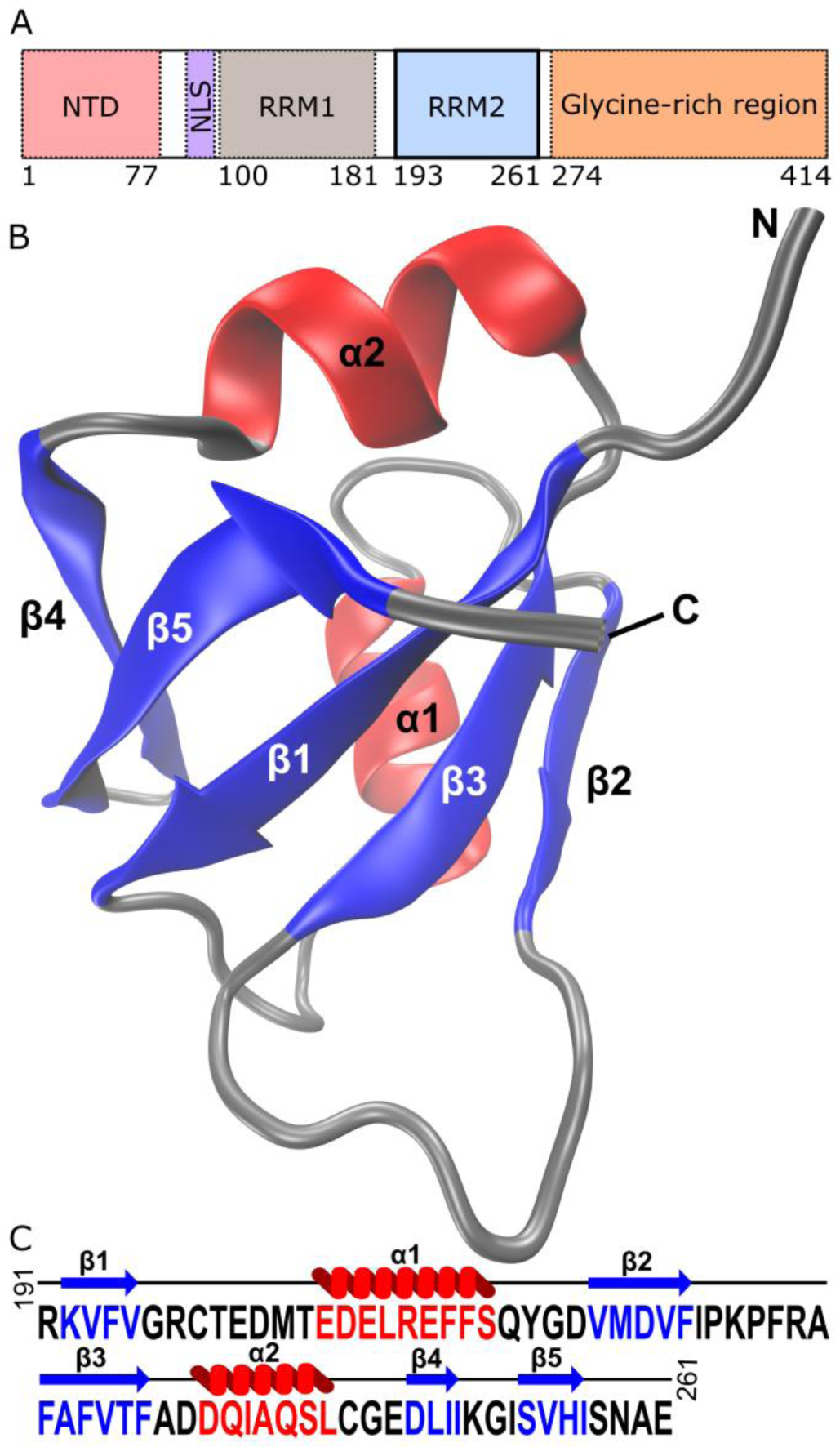
Domain architecture of TPD-43 with secondary structure and sequence of RRM2 domain. (A) Domain architecture of TDP-43 with terminal domain amino acid sequence numbers shown below. The RRM2 domain shown in blue comprises amino acids 191-261. (B) NMR structure (pdb: 1wf0) of the RRM2 backbone (gray ribbon) with β-strands shown in blue and α-helices shown in red. (C) Sequence of RRM2 with corresponding secondary structural elements shown above the sequence (β-strands in blue and α-helices in red).

While RRM1 has a high affinity for DNA and RNA (8), the function of RRM2 is less clear. RRM2 displays a relatively weak affinity for RNA (8) and is nonessential for the splicing ability of TDP-43 (9); however, RRM2 does enhance the specificity of RNA interactions (10). Structurally, RRM2 consists of a βαβ-repeat topology with an extra β-strand (β4, Fig. 1B, C) (8, 11). Unfolding studies have revealed that RRM2 is the stability core of TDP-43 and it populates a stable folding intermediate on its equilibrium unfolding pathway (12). While the precise roles for this folding intermediate in normal TDP-43 function and disease pathogenesis have yet to be determined, structural characterization would provide key insights into the potential mechanisms for decoupling function and dysfunction (13, 14).

In this study, the core nucleus of residual structure in the RRM2 folding intermediate was identified using a combined experimental and computational approach. The intermediate shows evidence of partial collapse with the residual secondary structure localized to the β_1_α_1_β_2_β_3_ region of RRM2, but it lacks a majority of the tertiary contacts observed in the native state. Using steered molecular dynamics (SMD) simulations and mutagenesis, we identified a critical interaction that maintains the structure of the RRM2 intermediate state. These findings can provide a framework for investigating the role of this folding intermediate within the stability core of TDP-43 in balancing proper folding and function with the risk of misfolding and disease.

## Results

### RRM2 remains partially collapsed with residual secondary structure in high urea

Low-resolution structural insights were obtained to probe the residual secondary and tertiary structure in the RRM2 intermediate state of TDP-43. Far-UV CD probed the changes in secondary structure upon unfolding of RRM2 in urea (Fig. 2A), where the unfolding profile reveals a three-state transition as shown in Scheme 1.

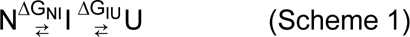

**Figure 2.**
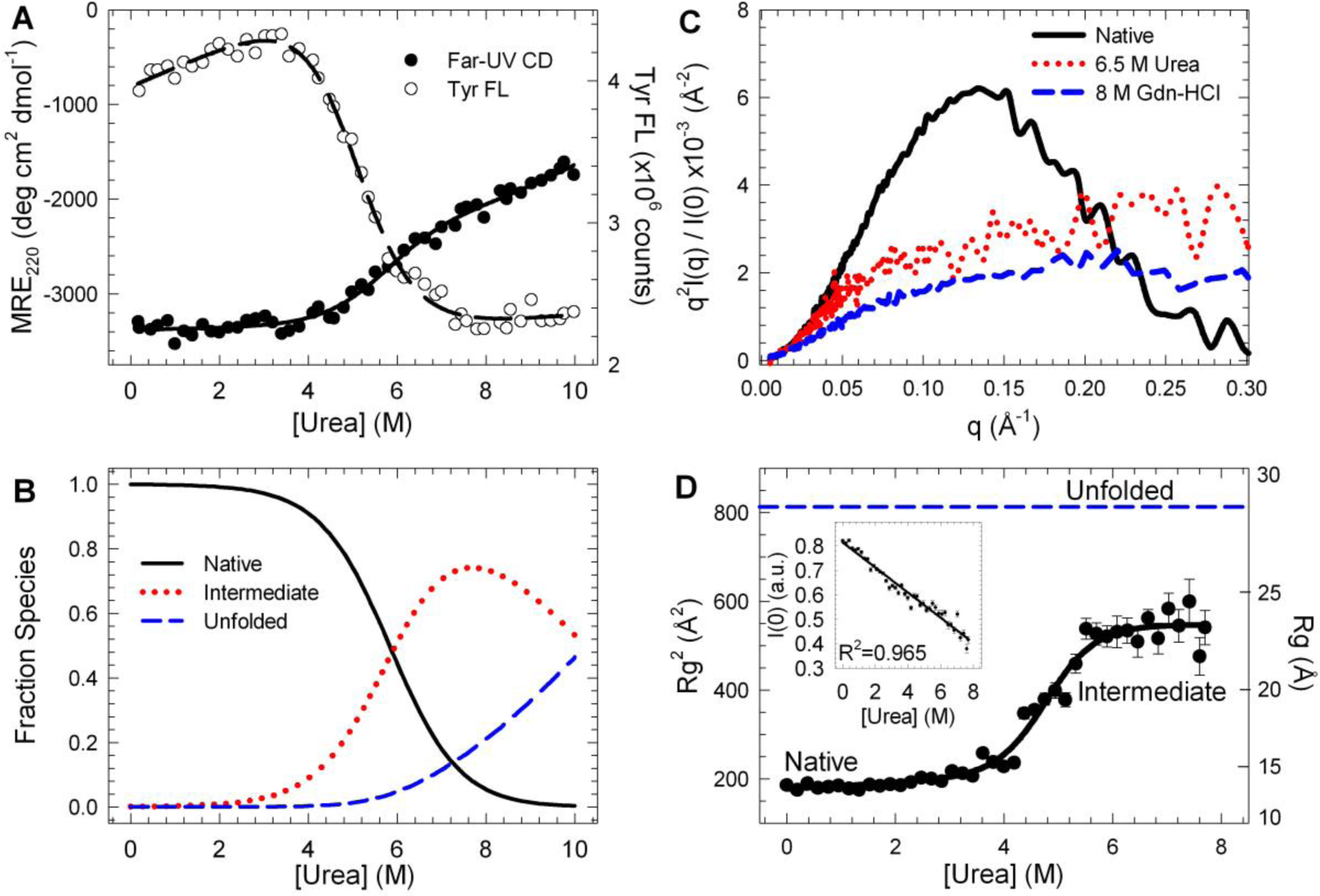
The RRM2 intermediate state lacks native globular structure but is not unfolded and has significant secondary structure. (A) Urea-induced equilibrium unfolding profiles of RRM2 monitored by far-UV CD (closed circles) and Tyr FL (open circles) were modeled to a three-state (NΔIΔU) and a two-state (NΔI/U) mechanism, respectively. (B) Population of the RRM2 native (solid), intermediate (red dotted) and unfolded (blue dashed) species as a function of urea obtained from the results of the fit of the far-UV CD. The intermediate state is 75% populated at 7.7 M urea. (C) Kratky plots of RRM2 under native conditions (black solid line), intermediate conditions at 6.5 M urea (red dotted line) and unfolded conditions at 8 M Gdn-HCl (blue dashed line). (D) Radius of gyration (Rg^2^) of RRM2 unfolding as a function of urea modeled to a two-state mechanism. The blue dashed line indicates the theoretical value for the unfolded state of RRM2. The plot of I(0) as a function of urea (inset) is linear (R^2^ = 0.965) as a function of denaturant, suggesting the increase in Rg^2^ is attributed to the intermediate state and not aggregation.

The N⇆I and I⇆U transitions contribute 4.30±0.10 kcal mol^-1^ and 3.38±0.13 kcal mol^-1^ to the overall stability of RRM2 (Table 1), in agreement with results obtained in Gdn-HCl (12). Using these thermodynamic parameters to determine the population of each species at a given concentration of urea (Fig. 2B), the RRM2 intermediate is the predominant species between 6.5 and 8.5 M, and never completely populates the unfolded state, even at high urea concentration (Fig. 2B; Fig. S1A). A singular value decomposition (SVD) analysis of the urea-induced unfolding profiles by far-UV CD revealed three component species that contribute to the observed CD spectrum. The reconstructed intermediate spectrum contains ∼50% of the native CD signal, indicative of residual secondary structure in the intermediate state (Fig. S1B).

**Table 1.** Thermodynamic parameters for the equilibrium unfolding of the RRM2 domain in urea. All experiments, except for far-UV CD experiments, were modeled to a two-state mechanism (N⇄I/U) to obtain the free energy of unfolding (ΔG°), m-value, and midpoint (C_m_) for the transition. In the case of the far-UV CD experiment, the data were modeled to a three-state mechanism (N⇄I⇄U). The m-value correlates to the buried surface area of RRM2, with the C_m_ representing the denaturant concentration where the two states are equally populated. Units: ΔG° (kcal mol-1), m (kcal mol-1 M-1) and C_m_ (M). Each experiment is the average of n≥3 experiments, except an average of 8 scattering profiles per urea concentration was used for SAXS. *The values reported by NMR are the weighted average of all secondary structural elements (all α-helices, β-strands and loops, with the exception of the N-terminal and first loops, where there were not enough data points). For the thermodynamic parameters of each element, see Table S2.

| | $\Delta G^\circ$ | m | $C_m$ |
| --- | --- | --- | --- |
| CD: N⇌I | $4.30 \pm 0.10$ | $0.73 \pm 0.02$ | $5.89 \pm 0.21$ |
| CD: I⇌U | $3.38 \pm 0.13$ | $0.33 \pm 0.01$ | $10.24 \pm 0.50$ |
| Tyr FL | $4.89 \pm 0.04$ | $0.92 \pm 0.10$ | $5.32 \pm 0.07$ |
| Near-UV CD | $7.29 \pm 0.33$ | $1.33 \pm 0.06$ | $5.48 \pm 0.35$ |
| SAXS | $6.26 \pm 1.09$ | $1.31 \pm 0.22$ | $4.78 \pm 1.16$ |
| *NMR | $5.00 \pm 0.96$ | $1.19 \pm 0.24$ | $4.22 \pm 0.24$ |

To further characterize this intermediate state, we used intrinsic tyrosine fluorescence (Tyr FL) and near-UV CD to probe the changes in the tertiary structure upon the population of this conformation. The Tyr FL unfolding curve yields a single cooperative transition (N⇆I/U) with a midpoint that is consistent with the N⇆I transition by far-UV CD (Fig. 2A), suggesting a lack of tertiary structure in the intermediate state. The native near-UV CD spectrum of RRM2 from 255 to 290 nm reveals a positive CD signal, resulting from the local packing of Phe and Tyr residues, which becomes drastically reduced when the intermediate is populated (Fig. S1C). The near-UV CD unfolding profile reveals a single cooperative transition (N⇆I/U) (Fig. S1D) with a midpoint that is consistent with the intrinsic Tyr FL unfolding profile (Table 1; Fig. S1E). In both cases, the expected total m-value is obtained in this single transition (Table 1), supporting the absence of defined tertiary structure in the RRM2 intermediate state.

Small-angle scattering (SAXS) measurements monitored changes in the size and globular structure of RRM2 with increasing urea. SEC-SAXS measurements of the native state reveal a monomeric species with an average radius of gyration (Rg) of 14.1±0.1 Å. The RRM2 domain is highly globular under these conditions (Fig. S2A), as supported by the parabolic shape of the Kratky plot compared to high urea and Gdn-HCl conditions (Fig. 2C). Rg measurements at increasing urea concentrations (Fig. 2D) reveal a single cooperative transition with a midpoint of 4.78±0.16 M (Table 1). The RRM2 intermediate state has an increased average Rg of 23.4±0.4 Å (Table S1) and reduced globularity (Figs. 2C and S2B) compared to the native state. The increased Rg at concentrations of >6 M urea is not a result of aggregation, as shown by the linear decrease of the intensity at zero scattering angle, I(0), with urea (Fig. 2D, inset) (15). As RRM2 does not completely unfold in urea (Fig. S1A), scattering as a function of Gdn-HCl concentration was used to investigate the unfolded state of RRM2 (Fig. S2C). At low concentrations of Gdn-HCl (<2 M), deviations from linearity of I(0) indicate aggregation is present (Fig. S2D), biasing the Rg toward larger species at those low Gdn-HCl concentrations. However, at concentrations >3 M, the intermediate state is highly populated (12) and has an Rg of 23.1±0.3 Å, similar to the Rg in urea (Table S1). At >6.5 M Gdn-HCl, the unfolded state dominates the unfolding equilibrium with an Rg of 28.5±1.2 Å (Table S1, Fig. S2C, blue), similar to the unfolded state of RRM1 (Fig. S2C, green). The theoretical Rg value (16) for the unfolded state of RRM2 is 27.7±2.6 Å (Fig. 2D, blue dashed line), larger than the observed Rg for the intermediate state (23.4±0.4 Å) but within error of the observed unfolded state for both RRM1 and RRM2. These results indicate that the RRM2 intermediate is not unfolded, consistent with the presence of residual secondary structural elements observed by far-UV CD (Fig. S1B).

### Residual secondary structure in the intermediate resides in the β_1_α_1_β_2_β_3_ region

NMR spectroscopy was used to characterize the structural differences between the RRM2 native and intermediate states with residue-level resolution (17, 18). The ^15^N-^1^H heteronuclear single-quantum coherence (HSQC) spectrum of the RRM2 native state (Fig. 3A, black) is well-dispersed in both the amide proton and nitrogen dimensions, as expected for a globular and structured domain (19). In comparison, the ^15^N-^1^H HSQC spectrum of RRM2 in 6.5 M urea (Fig. 3A, red), where the RRM2 intermediate state is the predominant species, is considerably more collapsed along both dimensions compared to the native state, but not as collapsed as a random-coil (20, 21). This result is consistent with a loss of the native globular structure observed by near-UV CD, Tyr FL and SAXS (Figs. 2 and S2) and the presence of residual secondary structure observed by far-UV CD (Figs. 2A and S1B) and SAXS (Figs. 2D and S2B).

**Figure 3.**
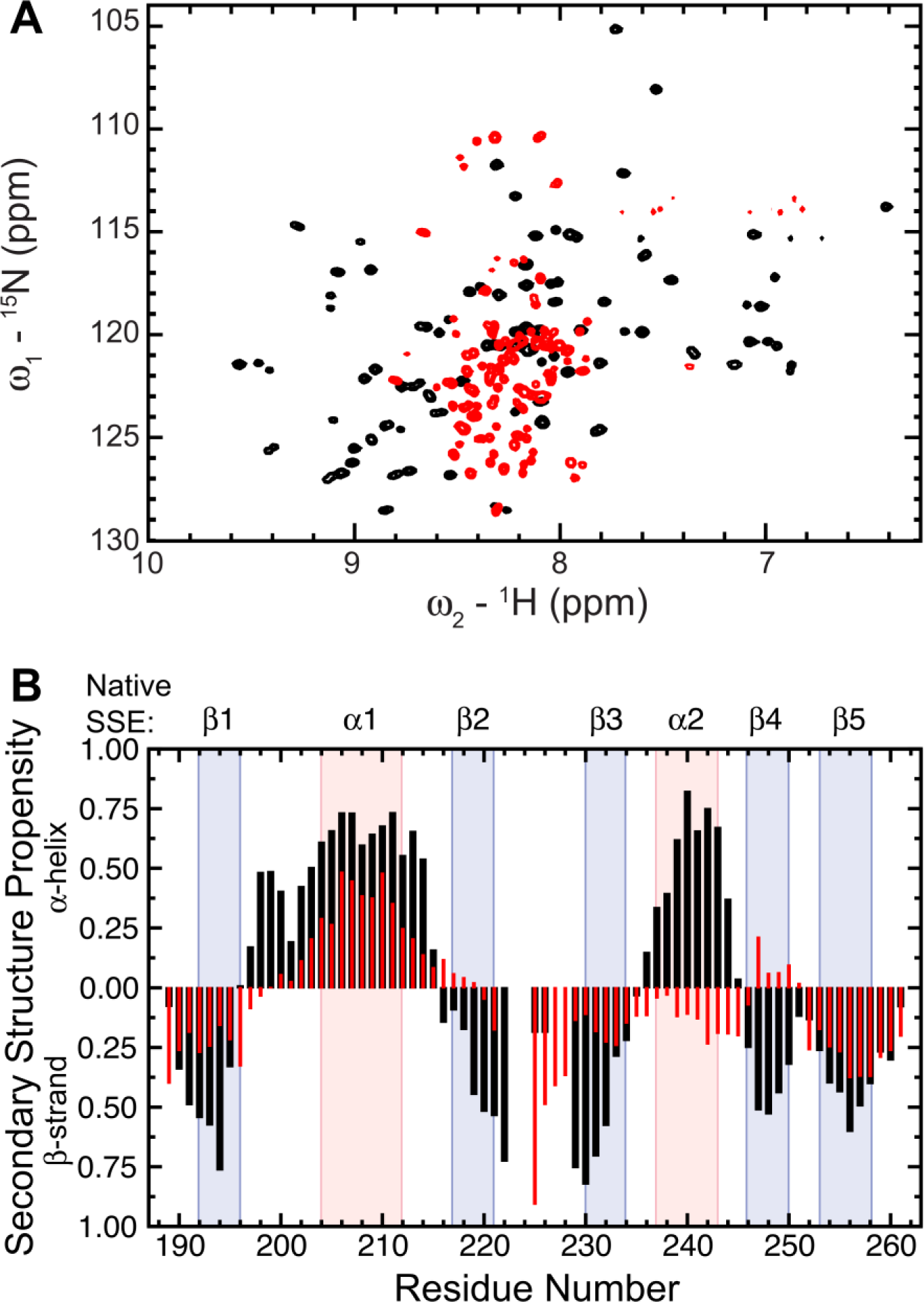
Residual structure in the intermediate state. (A) The ^15^N-^1^H HSQC NMR spectrum of the RRM2 intermediate state at 6.5 M urea (red) is collapsed in the ^1^H dimension compared to the native state (black). (B) Secondary structure propensity of the RRM2 native (black bars) and intermediate (red bars) states obtained from the ^1^H, ^15^N, Cα and Cβ chemical shifts deviations from random coil. Positive and negative values represent α-helical and β-strand propensity, respectively. Pink (α-helices) and light blue (β-strand) boxes highlight the secondary structural elements observed in the native structure.

The collected ^15^N-^1^H HSQC spectra between 0 M and 9 M urea show that different backbone amide groups are undergoing chemical exchange in different regimes (Fig. S3) (22). As the population of the RRM2 intermediate state increases with increasing denaturant concentration (Fig. S3A-E), we observed transitions in two different exchange regimes occurring at different urea concentrations in the HSQC spectra: 1) at urea concentrations lower than 6 M, a slow-to-intermediate chemical exchange regime dominates (cross-peaks from both states are observed per residue) and 2) at urea concentrations higher than 6 M, a fast-to-intermediate chemical exchange regime dominates (one cross-peak at the population average position). The appearance of fast and slow exchange characteristics suggests that the native and intermediate states are exchanging on a μs-ms intermediate timescale (23).

Chemical shift changes upon addition of urea are shown for several residues in Fig. S3A-E. Addition of urea under native favoring conditions (0-4 M urea: Fig. S3A-E, left) revealed that all residues within RRM2 behave differently depending on their location within the native structure. For example, V193 (β1) and I257 (β5) showed marked decreases in peak intensity at lower urea (∼2.0 M, yellow), while the peaks of E204 (α1), M218 (β2) and I250 (β4) persist until ∼4.5 M (green). At concentrations greater than 5.5 M urea, the peaks of the intermediate state dominate, increasing in intensity and experiencing additional chemical shift changes until the highest denaturant concentration tested (∼9 M urea) (Fig. S3A-E, right: burgundy to blue).

The native and intermediate states of RRM2 are exchanging in a slow-to-intermediate exchange regime at urea concentration < 6 M (22); therefore, it was necessary to assign the resonances of the intermediate state. Experiments to assign each residue’s backbone (^1^H, HN, Cα, CO) and Cβ were performed for both the RRM2 native and intermediate states as described in *Experimental Procedures*. The only unassigned residues in both states were those in the region 223-Pro-Lys-Pro-225.

Using the ^15^N-^1^H HSQC volumes of the cross-peaks corresponding to the native and intermediate states at each urea concentration, the population of the native state was plotted as a function of urea to determine the stability of secondary structural elements across the domain (Fig. S3F). Each residue within RRM2 undergoes a single cooperative transition that was modeled to a two-state unfolding mechanism (N⇆I/U) to obtain the thermodynamics of unfolding for each secondary structural element in RRM2 (Table S2). From this analysis we determined the ΔG, m-values (m) and transition midpoints (C_m_) for each secondary structural element and observed that the secondary structural elements that melt at the lowest concentration of urea are β5 and α2 followed by β3, β1, β2, β4 and α1 (Table S2). The average C_m_ of the exchange between the RRM2 native and unfolded states for all residues (C_m_=4.3±0.3 M) is lower than the midpoints obtained through near-UV, FL and SAXS measurements (Table 1). Taken together, these results suggest that the native and intermediate states may each consist of an ensemble of conformations that are spectroscopically similar by CD and FL, but distinguishable by NMR.

After assigning the backbone and Cβ chemical shifts of the native and intermediate states, the resonance frequencies were used to predict the amount of secondary structure using three different algorithms (SSP (20), SPARTA+ (24), and δ2D (25)) to identify residues that maintain a secondary structure propensity in the RRM2 intermediate state. Similar secondary structure predictions were obtained with the three methods (Fig. S4). In the case of the RRM2 native state (Fig. 3B, black bars), seven discreet regions deviate from random coil behavior: two with a high propensity for α-helices (α1: 203-211, α2: 237-244) and five with a high propensity for β-strands (β1: 190-195, β2: 216-222, β3: 229-234, β4: 246-250, β5: 254-260), in agreement with both the NMR structure (Fig. 1B) and DSSP (26). In 6.5 M urea (Fig. 3B, red bars), RRM2 has low propensity for secondary structure in the α2, β4 region, but preserves native-like propensity for secondary structure in β1 and α1. Two of the three algorithms predict loss of secondary structure in β5 (Fig. S4). Interestingly, the residues in the β2-β3 region continue to have β-strand propensity, albeit at loop residues between β2 and β3, suggesting a potential structural rearrangement. Overall, these results indicate that the N-terminus of RRM2 retains a native-like secondary structure propensity in the intermediate state, while most of the C-terminus, α2-β4-β5, adopts largely random-coil behavior.

### Mutations of the ILV cluster drastically alter the secondary structure of RRM2

Previous work has shown that clusters of buried, highly interconnected isoleucine, leucine and valine (ILV) residues are important for defining folding intermediates and cores of stability in globular proteins (27). RRM2 contains a single, large ILV cluster that comprises 16 residues with 19 contacts that span all secondary structural elements and is proposed to contribute to its enhanced stability compared to RRM1 and the formation of the intermediate state (12). Single alanine substitutions throughout the ILV cluster were designed to provide insights into the contribution of each residue to the stability and formation of the native and intermediate state of RRM2 (Fig. 4A).

**Figure 4.**
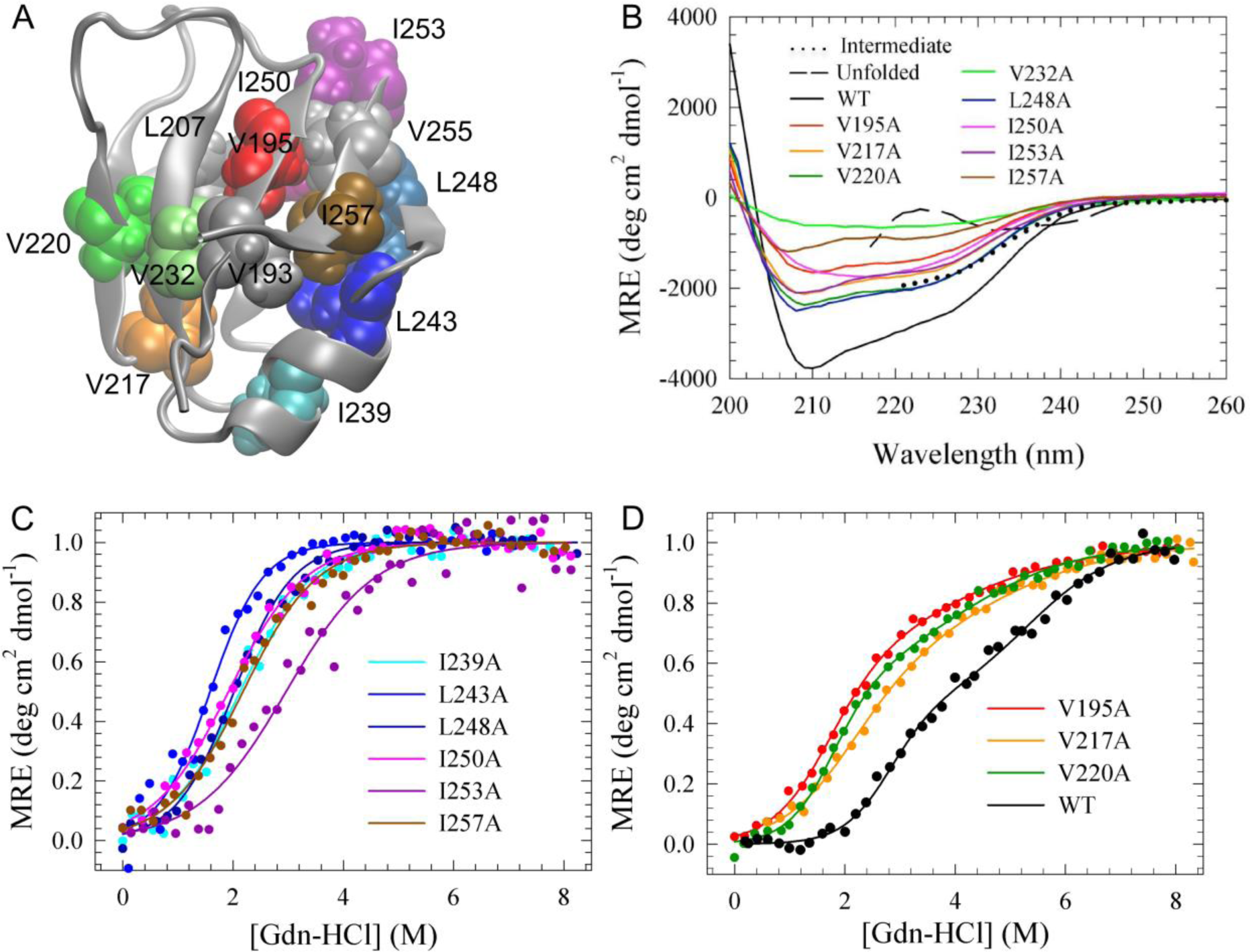
Alanine mutations in the ILV hydrophobic core drastically destabilize the native structure of RRM2. (A) Isoleucine, leucine and valine (ILV) residues that comprise the highly networked hydrophobic core mapped onto the RRM2 NMR structure (pdb: 1wf0) with appropriate residue labels. (B) CD spectra of the RRM2 mutants from 200-260 nm all show reduced secondary structure compared to the WT sequence (black), suggesting destabilization upon disruption of the ILV cluster. The RRM2 unfolded and SVD reconstructed intermediate states are shown as black dashed and dotted lines, respectively. (C, D) Fraction apparent plots of the CD equilibrium unfolding profiles of the RRM2 mutations fall into two separate classes. (C) Class I mutants were modeled to a two-state transition (N⇆U). (D) Class II mutants are destabilized relative to the WT (black) but retain a three-state equilibrium unfolding profile (N⇆I⇆U).

A native CD spectrum of each alanine variant, normalized for protein concentration, shows that all of the single alanine substitutions have drastically reduced MRE signals compared to the native RRM2 CD spectrum (Fig. 4B), suggesting an enhanced population of the intermediate state for the variants. While the overall shape of the WT RRM2 spectrum, characterized by a minimum at 210 nm and a shoulder at 230 nm, is maintained for most of the variants (Fig. 4B), the amplitudes are considerably reduced, ranging from 16-70% of the WT RRM2 MRE signal. Indeed, several of the variants examined resemble the reconstructed CD spectrum of the WT RRM2 intermediate (Fig. 4B), further supporting that the native states of these variants may resemble the RRM2 intermediate state.

Two variants, I250A (Fig. 4B, magenta) and V232A (Fig. 4B, light green), exhibit more pronounced effects. I250A displays a single minimum at 218 nm and lacks the 230 nm shoulder, indicating a significant alteration in the overall secondary structure of the domain rather than a mere reduction of the WT MRE signal. V232A, on the other hand, exhibits minimal secondary structure. These two variants appear to disrupt the overall secondary structure of the RRM2 domain, potentially leading to increased populations of the RRM2 intermediate or unfolded states.

Overall, the alanine scanning suggests that mutation of a single ILV residue of the hydrophobic core to alanine can have varying effects on the secondary structure of the isolated RRM2 domain that may result in increased populations of the RRM2 intermediate or unfolded states and may even, in some cases, yield RRM2 misfolded conformations.

### Alanine mutations enhance the population of the RRM2 intermediate state

The intriguing effect of the mutations on the secondary structure of RRM2 suggests the population of partially-folded and unfolded states are potentially enhanced by the removal of a single contact in the hydrophobic cluster. To test this possibility, the equilibrium unfolding profiles were obtained by monitoring the secondary and tertiary structures at increasing concentrations of Gdn-HCl using CD and Tyr FL (Fig. S5). Gdn-HCl was used as the denaturant instead of urea so that the unfolded form was fully sampled (Figs. 4C and 4D). While the Tyr FL behavior of the variants was quite variable, all of the variants exhibit a cooperative transition by CD, indicating a core of stability that is destabilized by denaturation (Figs. 4C and 4D). The equilibrium CD unfolding curve patterns for the variants were used to group the variants into two classes (Figs. 4C and 4D). Class I variants (I239A, L243A, L248A, I250A, I253A and I257A) exhibited unfolding profiles by CD that were modeled to a two-state mechanism (N⇆U) (Fig. 4C). The native-state CD spectrum of these variants matches more closely to the CD spectrum of the WT intermediate state, suggesting that their native structure is similar to the structure of the intermediate state of the WT in the absence of denaturant (i.e., N=I for these variants). Fraction apparent plots of the Class I mutations reveal similar midpoints for most of the variants, with the exception of L243A and I253A, suggesting a similar destabilization between these mutants (Fig. 4C). Each mutant variant is extremely destabilized compared to the WT with an average stability of ∼2 kcal mol^-1^ compared to the 7.42 ± 0.27 kcal mol^-1^ contributed from the N⇆I and I⇆U transitions observed for WT (Table S3). Furthermore, a majority of these mutations show a lack of denaturant dependence by Tyr FL (Fig. S5). By contrast, for WT RRM2, the Tyr FL unfolding partially correlates to the N⇆I transition, and the Tyr FL becomes insensitive to the I⇆U transition. The results suggest that the Class I mutants may lack the WT native tertiary structure as observed by the FL of Tyr214 and instead form a new “native” state, which may resemble the WT RRM2 intermediate state.

While some mutant variants result in a two-state unfolding profile, others, specifically V195A, V217A and V220A, retain an equilibrium unfolding profile with a population of the RRM2 intermediate and are referred to as Class II mutations (Fig. 4D). The V220A mutation most closely resembles the unfolding transition observed for the WT with destabilization of both the N⇆I and I⇆U transitions by 1.48 ± 0.16 kcal mol^-1^ and 1.91 ± 0.18 kcal mol^-1^, respectively. Similarly, the midpoint of the Tyr FL (1.89 ± 0.04 M) coincides with the N⇆I transition by CD. Each transition of the unfolding profile of V195A and V217A is further destabilized compared to V220A but still provides ∼1.8 kcal mol^-1^ from each transition (Table S4). The fraction apparent unfolded plots of the Class II mutations, in comparison to WT, reveal shifted midpoints toward lower denaturant concentrations (Fig 4D), suggesting an increase in the population of the intermediate state. The thermodynamic parameters obtained from the fitting of the three-state model confirm this increase and show that all of the Class II mutations increase the population of the RRM2 intermediate state by an order of magnitude (10-25 fold) (Table S4, I (%)).

### The V193-V232 hydrophobic contact is critical to the stable core of RRM2

The V232A variant is unique and does not fall into either Class I or Class II as discussed above. Instead, this variant exhibits relatively little secondary structure (Fig. 4B) and no apparent denaturant dependence, suggesting that this residue may be critical for maintaining the core of structure in the WT RRM2. To explore the core of stability further, constant force steered molecular dynamics (SMD) was performed as a complimentary method to investigate the relative stability of the secondary structural elements of RRM2 and provide atomistic details of the unfolding process (28). SMD has successfully identified folding intermediates (29) and has also augmented experimental mechanical unfolding studies (30).

The extension profiles of the RRM2 unfolding over a range of constant forces are shown in Fig. 6 (left), with the representative structure from each conformational state observed at different simulation times shown as Roman numerals for the extension profile at a constant force of 225 pN (red). As shown in Fig. 6, the order of secondary structure unfolding, from least to most stable, is β4/β5 (II), α2 (III), β1 (IV), α1 (V), and β2/β3 (VI); this order is insensitive to initial conditions and pulling from either the N- or C-terminus (Fig. S6). All extension profiles of RRM2 that undergo unfolding (forces ≥200 pN) exhibit a plateau at ∼90 Å, indicating the presence of a stable intermediate (Fig. 6, conformation III) before complete unfolding at longer timescales (Fig. 6, conformations IV-VI). Conformation III retains secondary structure in β1, α1, β2 and β3 (Fig. 6), consistent with the secondary structure revealed by NMR at 6.5 M urea (Fig. 3B) and by the urea titration data obtained by NMR (Table S2). Therefore, the N-terminus provides a core of stability in both the mechanical- and denaturant-induced unfolding of RRM2 (Fig. 6, conformation III).

**Figure 5.**
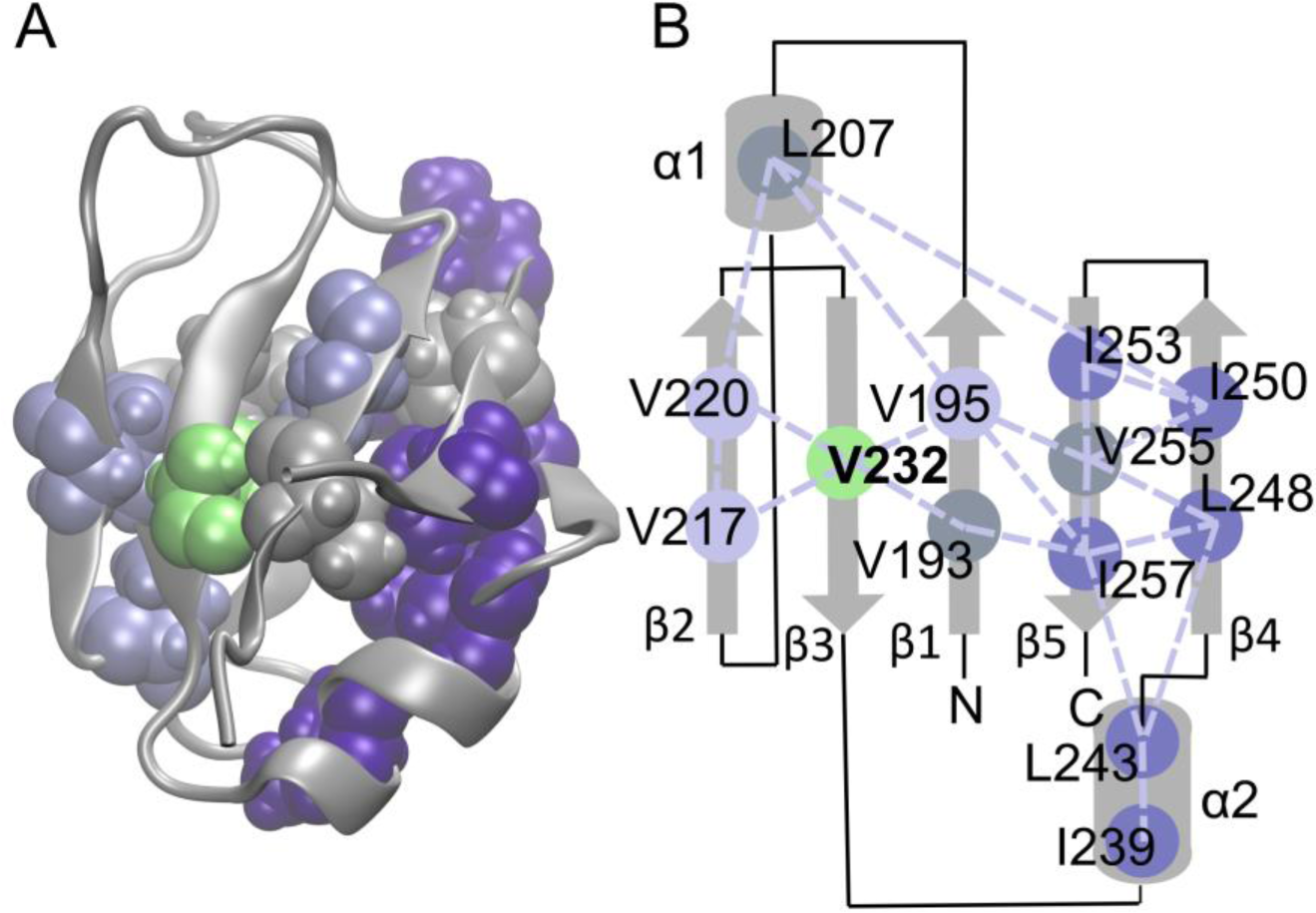
Alanine mutations alter the folding energy landscape of RRM2. ILV residues, shown as spheres, mapped onto (A) the NMR structure (pdb code: 1wf0) and (B) the topology map of RRM2. Dashed lines indicate hydrophobic contacts identified from the ensemble of NMR structures. The highly destabilized Class I mutations (dark purple) cluster in the C-terminal half of RRM2, while the more stable Class II mutations (light purple), which exhibit three-state behavior, are localized to the N-terminus. V232, which is highly destabilizing when mutated, is shown in green. Residues in gray were not mutated to alanine.

**Figure 6.**
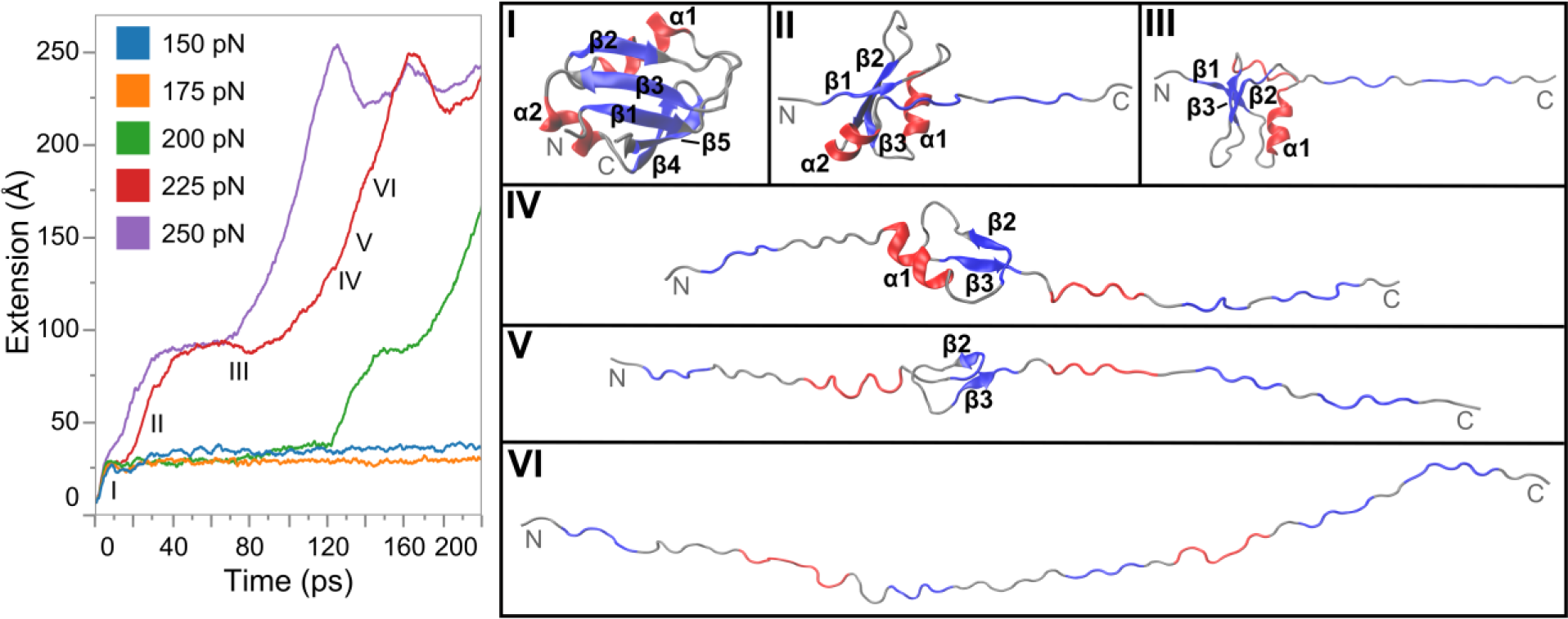
The N-terminal half of RRM2 is the core of stability in the intermediate state. Extension profiles (left) as a function of time from constant force steered molecular dynamics of RRM2. Roman numerals (I-VI) mark the locations of the conformational states of the unfolding pathway under 225 pN of force (red). A representative structure from each state is shown on the right. Gray ribbon shows RRM2 backbone with β-strands in blue and α-helices in red. The secondary structural elements maintained in each representative structure are labeled. The plateau at 90 Å indicates the existence of an intermediate state (conformation III).

A residue-level analysis of the SMD results provides molecular details into the specific side chain contacts that stabilize the RRM2 intermediate state and shed light on why some alanine mutations in the hydrophobic core are more disruptive to the structure of RRM2. The loss of the hydrophobic contacts between the ILV residues in RRM2 was correlated with conformations I-VI observed in the extension profiles (Fig. S7), showing an average of 8-9 ILV contacts in the intermediate state (conformation III). The observed hydrophobic contacts as a function of pulling time (Fig. S8) shows specific ILV contacts that are most important for maintaining the different partially-unfolded structures on the unfolding pathway. The unfolding of the C-terminus occurs rapidly with the loss of the hydrophobic contacts between α2, β4, and β5 (Fig. S8, red and orange). I250, a highly disruptive Class I mutation (Fig. 4, Table S3), is located adjacent to the turn between β4 and β5 – loss of the L207-I250 contact is the final part of the unravelling of β4 and β5 (Fig. S8, red), suggesting a more critical role in stabilizing these secondary structural elements. L243, the most destabilizing Class I mutation (Fig. 4, Table S3), is the major hydrophobic contact broken with the loss of α2 (Fig. S8, orange). Once the contact between L243 and I239 is broken, the protein becomes more extended and the conformational population switches from III to IV (Fig. 6). The computational results can also explain the difference in stability shown by the I239 and L243 mutations (Fig. 4, Table S3). I239 is the less disruptive mutation in this contact pair because it only contacts L243. By contrast, the L243 mutation is more disruptive because it has additional transient contacts with ILV residues in β1, β3, and α1 (Fig. S8).

The plateau in the extension profile following these unfolding events (Fig. 6, III) corresponds to the reorientation of β1/β3 from anti-parallel to a perpendicular configuration, and this structure is maintained by the hydrophobic contact between V193 (β1) and V232 (β3) (Fig. 7). Conformations IV-VI occur only when this contact is broken (Fig. S8, green), implicating the critical importance of the V193/V232 contact in maintaining the stable core of RRM2. This importance is supported by the alanine scanning: only the V232A mutation severely destabilizes the native state and results in a significant population of the RRM2 unfolded state under native conditions (Fig. 4B). An alternative approach using replica-averaged metadynamics based on the experimental chemical shifts at 6 M urea (31) identified multiple partially folded states, several of which maintain the same secondary structural elements observed in the SMD conformations. The metadynamics results similarly identified the importance of the V193/V232 contact in maintaining the more structured microstates. Combined, these results demonstrate that V232 is a critical keystone residue that cements the residual structure in the RRM2 intermediate state and also maintains the overall fold of this RRM domain in TDP-43.

**Figure 7.**
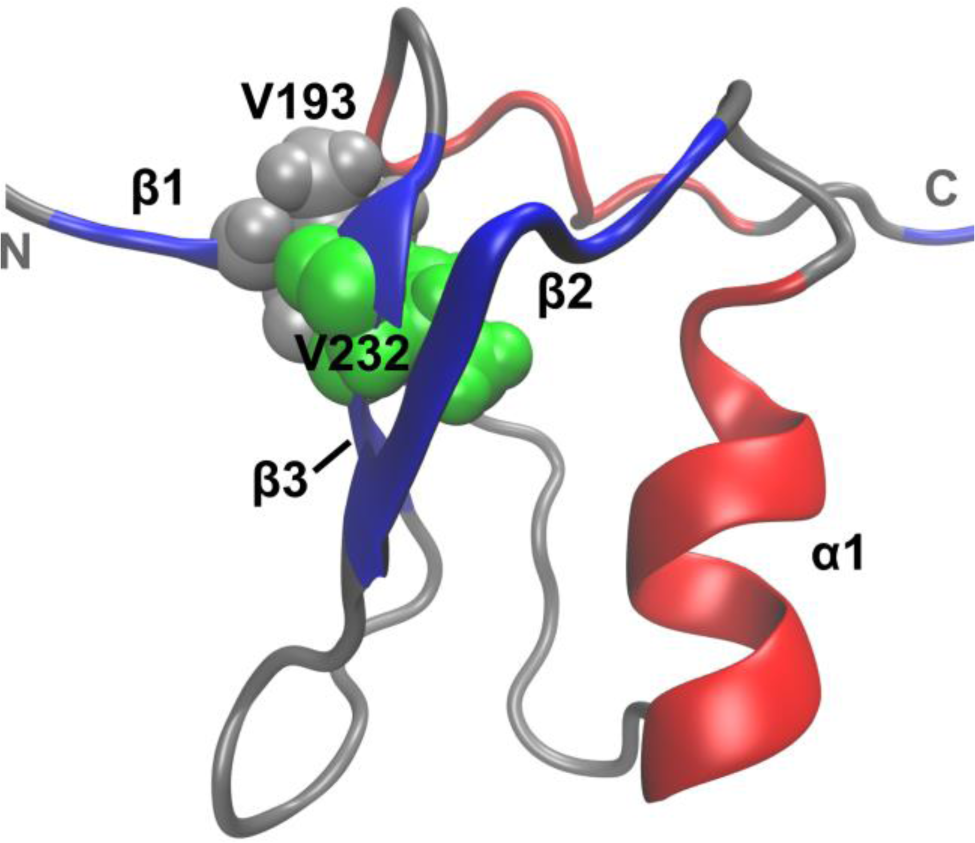
V232 plays a key role in stabilizing the residual structure in the intermediate and native states of RRM2. Core residual structure in the RRM2 intermediate state, where the V193/V232 (β1/β3) hydrophobic contact prevents further unfolding. The van der Waals representations of V193 (gray) and V232 (green) show the hydrophobic contact on the gray RRM2 backbone of the maintained structure in conformation III with β-strands in blue and α-helices in red.

### N-terminus provides a core of stability during unfolding

Combining the insights derived from NMR spectroscopy, alanine scanning and SMD allows us to hypothesize about the difference between (and existence of) Class I and Class II mutations. Class I mutations are located in the secondary structural elements that are lost first in the extension profiles (β4, β5, and α2, Figs. 5 and 6) and lack secondary structure at 6.5 M urea according to NMR (Fig. 3B). The absence of structure in these secondary structural elements in the intermediate is consistent with a two-state nature for these mutations: these mutations destabilize β4, β5 and/or α2, resulting in a native state that resembles an intermediate-like structure. Thus, there can be no native-to-intermediate transition in the unfolding profiles and the data therefore fit to a two-state model.

The mutations that fit instead to a three-state model (Class II) are located in the most stable secondary structural elements. These elements are lost after the plateau in the extension profile (β1, β2, β3, and α1, Figs. 5 and 6) and were observed to maintain some secondary structure at 6.5 M urea (Fig. 3B). Therefore, these secondary structural elements are less likely to be fully disrupted by a single point mutation. Since the less stable secondary structural elements of β4, β5, and α2 are not directly perturbed by these mutations, they are likely to maintain a native-like fold, and therefore upon addition of denaturant could transition from a native-like fold to an intermediate-like structure where β4, β5, and α2 are lost. Therefore, even though the mutant native states do not fully resemble the WT native state, Class II mutations result in an apparent three-state model presumably because the stable hydrophobic core remains significantly more stable than β4, β5, and α2.

The SMD results suggest that there may be two different mechanisms that lead to an increase in the population of the intermediate state in Class II mutations. In the case of V195A, this residue forms hydrophobic contacts with β4 and β5 (Fig. S8); loss of this hydrophobic residue could destabilize these secondary structural elements, increasing the population of the intermediate state. For V217A and V220A, the increased population of the intermediate state could indicate that although there is a stable core comprised of β1, β2, β3, and α1, the conformation of this core may differ between the intermediate state and the native state, consistent with the reorientation of β1/β3 we see in the extension profile plateau (Figs. 6 and 7) and the altered β-strand propensity observed for β2/β3 in the NMR results at 6.5 M urea (Fig. 3B). Mutation of these residues to alanine could facilitate a conformational transition to a more intermediate-like structure in the stable core.

The V232A mutation is catastrophic to the secondary structure of RRM2, making this mutant the most destabilized of those tested in this study (Fig. 4B). Due to its almost complete lack of secondary structure, this mutant failed to fit to a two-state or three-state model and is not likely to populate conformations resembling either the native or intermediate structures of WT RRM2 (Fig. 4B, green).

Together, the results of this study suggest the following characteristics for the intermediate state: 1) loss of structure of β4, β5, and α2, 2) a stable hydrophobic core of residual structure comprised of β1, β2, β3, and α1 that does not have an entirely native-like conformation, and 3) a critical role for V232 in the stability of this hydrophobic core.

## Discussion

We used a combined experimental and computational approach to investigate the structural characteristics of the stable folding intermediate of RRM2 in the ALS-linked protein, TDP-43. This intermediate state lacks the globular structure and tertiary contacts of the native state (Fig. 2) but contains significant secondary structure in the N-terminal half of RRM2 (Fig. 3B). In contrast, the C-terminal half of RRM2 appears more disordered (Fig. 3B) and unfolds first during pulling simulations (Fig. 6). Furthermore, the hydrophobic interaction between V193 and V232 is critical for the formation of the intermediate and native states (Figs. 4 and 7). The lower stability of the RRM2 C-terminus could result in the exposure of normally buried hydrophobic residues and aggregation-prone sequences (32, 33) to solvent, providing a binding surface for functional intra-and intermolecular interactions or possibly for misfolding and aggregation with other TDP-43 molecules and similar proteins (34).

RNA binding plays a critical role in the functions of TDP-43 as well as its localization in the nucleus (35, 36). Previous work (12) has demonstrated that interactions with RRM1 and/or RNA stabilize the native state of RRM2 against the population of the intermediate state. Indeed, the NMR structure of the tethered domains shows hydrophobic contacts between the C-terminal domain of RRM2 and β2 and β3 of RRM1 (37). The RRM2 intermediate may also serve a crucial functional role in nucleocytoplasmic transport of TDP-43 in several ways. First, partial unfolding of RRM2 impacts RNA binding, shifting the localization from nuclear to cytoplasmic (35-37). Second, increased concentration of TDP-43 in the cytoplasm may affect stress granule formation and the formation of insoluble aggregates (38). Finally, exposure of hydrophobic residues in the RRM2 intermediate may also affect interactions with chaperones such as with HSP70 (39), altering overall protein homeostasis (40).

Cleavage of TDP-43 has been observed in patient samples at multiple cleavage sites (41) and can result in the degradation of the N-terminal region, including RRM1. This mechanism results in the accumulation of C-terminal fragments comprised of RRM2 and the C-terminus that are toxic and highly aggregation prone (7). While the role of cleavage in disease pathogenesis is highly controversial (2, 4, 5), a disease-relevant fragment of RRM2 shows increased populations of the intermediate state (12) and increased aggregation propensity (32).

Our study reveals a loss of tertiary structure and globularity of RRM2 upon the population of the intermediate state (Fig. 2). Multiple disordered proteins linked to neurodegeneration can undergo large conformational changes (42) that are thought to be more aggregation-prone than their fully disordered counterparts and could result in the formation and propagation of aggregates and fibrils (43). Two examples are the peptide Aβ and α-synuclein. While Aβ appears completely disordered and α-synuclein has some helical propensity, both transiently sample pathogenic conformations (44-46). In a similar manner, the RRM2 intermediate may sample alternative conformations that interact aberrantly with the adjacent intrinsically disordered, prion-like glycine-rich C-terminal domain (47). Thus, the RRM2 intermediate, which is sampled at low frequency under native conditions (12), may exert toxicity through comparable mechanisms as other disease-related proteins (48, 49). The mutant variants of RRM2 presented in this work (Fig. 4), which preferentially populate the intermediate species, if placed in the context of full-length TDP-43, can serve as tools to decipher the molecular basis of disease. In addition, they can provide a target for antibody and biomarker development for use in the diagnosis and treatment of TDP-43-mediated neurodegenerative disease (50, 51).

## Experimental procedures

### Protein Expression and Purification

RRM2 was expressed and purified as described previously (12), supplemented with a size exclusion chromatography step (Superdex75, GE Healthcare) in the experimental buffer (20 mM MOPS pH 6.8, 25 mM KCl and 1 mM β-mercaptoethanol). For ^15^N and ^13^C protein samples, M9 minimal media containing 1 g L^-1^ of ^15^N-NH_4_Cl and 2 g L^-1^ ^13^C-glucose was used for protein expression. The protein purity was >98% as determined by SDS-PAGE and MALDI-TOF mass spectrometry carried out at the Proteomics and Mass Spectrometry Facility at the University of Massachusetts Medical School. Protein concentration was measured by A_280_ absorbance using an extinction coefficient of 1490 M^-1^ cm^-1^.

### Size Exclusion Chromatography coupled with Small Angle X-ray Scattering (SEC-SAXS)

Native state SEC-SAXS measurements were acquired at the BioCAT beamline at the Advanced Photon Source (APS), Argonne National Laboratory (Argonne, IL, USA). Scattering images of monomeric RRM2 were obtained using an in-line liquid chromatography system prior to X-ray exposure to remove larger aggregated species, if present. RRM2 (>5 mg mL^-1^) was injected onto a 24 mL Superdex 75 size exclusion column (GE Healthcare) at a flow rate of 1 mL min^-1^. Images were acquired using a Mar165 CCD detector (MarUSA, Inc., Evanston, IL) with ∼4 sec intervals between ∼2 sec exposures. The buffer-corrected SAXS images of the SEC elution profile were analyzed using Igor Pro (Wavemetrics, Inc., Lake Oswego, OR) to obtain the concentration dependence, radius of gyration (Rg) and pairwise distribution functions, which were required to determine the average electron density envelopes using ATSAS (32). The NMR structure of RRM2 (pdb: 1wf0) was modeled into the electron density envelope using Chimera (52).

### Equilibrium Unfolding Experiments

All far-UV CD and Tyr FL equilibrium unfolding experiments were performed at 25 °C as described previously (12). For near-UV experiments, spectra were collected with 0.5 mg mL^-1^ RRM2 using a 5 cm cuvette (Hellma) and averaged over twenty scans. SAXS measurements were acquired for ∼1.5 mg mL^-1^ RRM2 at increasing urea and Gdn-HCl concentrations as described above, with the exceptions that the in-line size exclusion chromatography set up was not used for the urea titration and the exposure time was ∼1 sec at each urea concentration. The radius of gyration (Rg) and the intensity at zero scattering angle (I(0)) at each urea concentration was determined by the Guinier approximation in Igor Pro (Wavemetrics, Inc., Lake Oswego, OR). The deviation of the Rg from the theoretical value for an unfolded random coil peptide was calculated using the Rg dependence on amino acid length (16).

Two-dimensional ^15^N-^1^H HSQC spectra at increasing denaturant concentrations were obtained using a Varian 600 MHz spectrometer at 25°C using 1.5 mg mL^-1^ RRM2 in 92% H_2_O / 8% D_2_O over 64 scans. The fraction native for each residue was determined from the peak volumes at each concentration of urea for the native and intermediate state assignments using Equation 1:

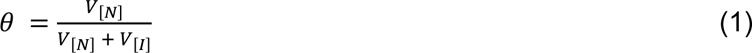

where θ is the fraction native and V_[N]_ and V_[I]_ are the volumes of the native and intermediate cross peaks, respectively, at a given concentration of urea. The fraction natives were averaged for each secondary structural element and used to determine ΔG, m-values (m) and transition midpoints (C_m_) Table S2). The chemical shift perturbation (CSP) with increasing denaturant concentration was determined using the native spectrum of RRM2 as a reference (53).

All urea denaturation experiments, including near-UV, Tyr FL, SAXS and NMR measurements, were modeled to a two-state thermodynamic transition, N⇆I/U (12), as the unfolded state is not significantly populated at high urea concentrations (>9 M). The I(0) values as a function of denaturant from the Guinier analysis of the SAXS profiles were modeled using a linear regression. The far-UV CD urea denaturation unfolding profile was modeled to a three-state transition using the baseline and slope of the unfolding baseline of RRM1 at high urea, as the unfolding baselines and slopes of RRM1 and RRM2 have been previously shown to be similar under high concentrations of Gdn-HCl (12). All reported free energies in the absence of denaturant (ΔG°), m-values (m) and transition midpoints (C_m_) are the global fit to three independent experiments. Singular value decomposition (SVD) analysis of the far-UV CD spectra as a function of denaturant to obtain the component spectra of the native, intermediate and unfolded states of RRM2 in urea and Gdn-HCl was performed as previously described (54). Populations of species of the RRM2 equilibrium unfolding profile were determined using the global fit of the far-UV CD data (12).

### NMR Assignment and Residual Structure Determination

Triple resonance assignment experiments were performed using a ^13^C,^15^N-labeled 4 mg mL^-1^ RRM2 sample in the native (0 M urea) and intermediate (6 M urea) states (BMRB deposition number 27549). Chemical shifts for each state’s backbone (HN, N, Cα) and side chains (Cβ) were obtained through HSQC, HNCA, HN(CO)CA, CBCA(CO)NH, HNCACB experiments. The native state was further supplemented with carbonyl (CO) assignments using HNCO and HN(CA)CO experiments (53).The spectra were processed using NMRPipe (55) and assigned and analyzed using Sparky (T. D. Goddard and D. G. Kneller, SPARKY 3, University of California, San Francisco). Secondary structure propensity in the RRM2 native and intermediate states were determined using the appropriate chemical shifts as described by Marsh *et al* (20), SPARTA+ (24) and δD2 (25).

### Steered Molecular Dynamics

The simulations used residues 189 to 261 of the NMR solution structure of TDP-43 RRM2 (pdb: 1wf0) with the wild-type loop residue mutations S191R, G192K, and G200E. Three different equilibration procedures were used to obtain the initial conditions for the SMD simulations: explicit solvent, implicit solvent, or implicit solvent with the Linear Combination of Pairwise Overlap method (LCPO) for adding the nonpolar/hydrophobic energy contribution (56).

The NMR structure was minimized and equilibrated in explicit solvent at 295 K with neutralizing counter ions according to the protocols reported in our previous work (57). Three independent, equilibrated structures collected after 20 ns from these explicit solvent simulations were used as the initial conditions for the SMD calculations.

Implicit solvent simulations were performed in NAMD 2.10 (58) with the generalized Born implicit solvent model. A temperature of 298 K was maintained by Langevin dynamics with a Langevin damping coefficient of 50/ps and the ionic concentration was chosen to match the neutralized explicit solvent simulations (0.070 M). Three independent trajectories were collected in implicit solvent both with and without the LCPO method using the following protocol. The NMR structure was minimized using the conjugate gradient method with all backbone atoms fixed followed by a second conjugate gradient minimization with the position of the Cα atoms constrained with harmonic restraints. The initial scaling of the constraint force by 5 was slowly decreased to zero (no constraints) over 0.5 ns followed by 10 ns of unrestrained structural equilibration to generate the initial conditions for the SMD calculations.

Implicit solvent simulations of RRM2 without the use of the LCPO method for adding the nonpolar/hydrophobic energy contribution from the implicit solvent were found to have a solvent accessible surface area (SASA) 10% greater than the SASA of the NMR structure. The LCPO method with a value of 0.11 kcal/mol/Å^2^ for the surface tension was found to reproduce the SASA of RRM2 seen in the NMR structure and explicit solvent simulations (Figure S9A). An additional 240 ns were collected in implicit solvent using LCPO to confirm the consistency of results with the published structure and between solvent models (Figure S9).

Steered molecular dynamics were performed under a constant force using NAMD 2.10 (58). The Cα of the N-terminal residue (G189) was fixed while a force was applied to the Cα of the C-terminal residue (E261). SMD was performed using three initial configurations generated by the three different equilibration procedures as mentioned above. Each initial configuration was pulled for 200 ps with constant forces of 100, 150, 175, 200, 225, 250, 275, 300, 400 and 500 pN. The order of secondary structural element unfolding was found to be insensitive to initial structure and velocities. The initial conditions for the representative unfolding pathway presented in this work were obtained from equilibration in implicit solvent with LCPO. As an additional validation, the SMD calculations were repeated for the initial conditions presented in this work with the fixed and pulling atoms reversed (G189 pulled, E261 fixed). The order of unfolding and the secondary structural elements preserved in the predicted intermediate were not altered by reversing the pulling direction (Fig. S6).

### Analysis of Trajectories

VMD 1.9.2 (59) was used to process the trajectories and calculate the metrics described below, and to visualize the structures using the STRIDE method (60). The data generated was processed and plotted using Tableau Software v8.3 (Seattle, WA).

### Hydrophobic contacts between ILV residues

Hydrophobic contacts were defined by the amount of surface area buried between a pair of residues using Equation 2:

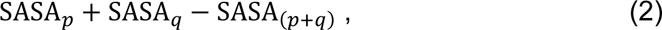

where p and q are ILV residues and SASA is the solvent accessible surface area of the side chain of residue p, q, and residues p and q together. This calculation determines the amount of surface area buried by the contact between residues p and q. The hydrophobic contact probability is one if two residues have at least 45 Å^2^ of buried surface area and zero otherwise. A cutoff area of 45 Å^2^ was chosen based on previous work (12, 27, 61, 62).

### Extension profiles

The extension profiles of the SMD simulations were calculated as the distance between the Cα of G189 and the Cα of E261 as a function of time.

### Mutational Analysis

Mutations in the hydrophobic isoleucine, leucine and valine (ILV) clusters of the isolated RRM2 domain were guided by an in-house software that calculates the contact surface area between residue side chains (27). ILV to alanine (A) mutations were inserted into the WT isolated (residues 193 – 261) RRM2 domain in a modified pGEX-6p1 vector (4) using the QuickChange method (Stratagene) and verified through Sanger Sequencing (Genewiz). All constructs have an N-terminal hexahistidine (His_6_) tag with a PreScission protease cleavage site (LEVLFQ/GP) to aid in protein isolation.

All mutant RRM2 proteins were transformed into BL21 DE3 PlysS *E. coli* for protein expression after induction at 30°C by 1 mM Isopropyl β-D-1-thiogalactopyranoside (IPTG) for 16-20 hours. Purified protein was isolated as previously described (4) using metal-affinity chromatography followed by PreScission protease digestion and Q sepharose ion exchange chromatography. The purity of all proteins was >95 % as indicated by SDS-PAGE. Protein concentration was measured by A_280_ absorbance (41), using an extinction coefficient of 1490 M^-1^ cm^-1^ for RRM2 mutant constructs.

The native-state circular dichroism (CD) spectrum of each mutant in the isolated RRM2 construct was collected from 200-280 nm on a Jasco-810 spectropolarimeter with a thermoelectric temperature control system in a 0.1 cm cuvette (Hellma). Guanidine hydrochloride (Gdn-HCl)-induced denaturation spectra were collected from 260-215 nm at a scan rate of 50 nm min^-1^ and a response time of 8 s for each isolated RRM2 mutant. Samples were prepared as previously described (4). All Gdn-HCl concentrations were measured using an ABBE Refractometer, and all CD measurements were baseline corrected for buffer contributions. Each CD spectra was normalized for protein concentration and number of amino acids and reported as mean residue ellipticity (MRE) (63). Steady-state fluorescence (FL) measurements were performed on a Spex Fluorolog-3 equipped with a wavelength electronics temperature controller for the isolated RRM2 constructs as previously described (4).

Denaturation experiments by CD and FL were performed in triplicate for each mutant. The equilibrium folding data were analyzed using an appropriate equilibrium folding model with the in-house data analysis software Savuka (64) as previously described. Each data set was subjected to a global analysis, where the baselines were local parameters and the free energy of folding in the absence of denaturant (⇆G°_H2O_) and the *m* value were globally linked between data sets.

## Data availability

The backbone chemical shift assignments for TDP-43 RRM2 at 6M urea have been deposited with accession code BMRB: 27549. The raw MD trajectories are available upon reasonable request. All other data are contained in the manuscript and supporting information.

## Supporting information

This article contains supporting information.

## Conflict of interest

The authors declare that they have no conflicts of interest with the contents of this article.

## Acknowledgments

We are grateful to Osman Bilsel, Daryl Bosco, Srinivas Chakravarthy, Tom Irving, Bob Matthews, Sean Ryder and Zuoshang Xu for insightful discussions.

## Funding and additional information

This work was supported by grants from the ALS Association (16-11P-280), the ALS Therapy Alliance, Inc. and the National Institutes of Health (GM137529, GM54836, GM103622 and RR008640). This research used resources of the Advanced Photon Source, a U.S. Department of Energy (DOE) Office of Science User Facility operated for the DOE Office of Science by Argonne National Laboratory under Contract No. DE-AC02-06CH11357. BioCAT was supported by grant P41 GM103622 from the National Institute of General Medical Sciences of the National Institutes of Health. The content is solely the responsibility of the authors and does not necessarily reflect the official views of the National Institute of General Medical Sciences or National Institutes of Health.

## Abbreviations

ALS: Amyotrophic Lateral Sclerosis
CD: Circular Dichroism
DSSP: Define Secondary Structure of Proteins
Gdn-HCl: guanidinium hydrochloride
HSQC: Heteronuclear Single-Quantum Coherence
ILV: Isoleucine, Leucine, and Valine
LCPO: Linear Combination of Pairwise Overlap
NMR: Nuclear Magnetic Resonance
Rg: Radius of gyration
RRM: RNA Recognition Motif
SASA: Solvent Accessible Surface Area
SAXS: Small Angle X-ray Scattering
SEC-SAXS: Size-Exclusion Chromatography Small Angle X-ray Scattering
SMD: Steered Molecular Dynamics
SVD: Singular Value Decomposition
Tyr FL: Tyrosine Fluorescence

## Notes

### Competing Interest Statement

The authors have declared no competing interest.

## References

1. Sreedharan, J., Blair, I. P., Tripathi, V. B., Hu, X., Vance, C., Rogelj, B. et al. (2008) TDP-43 mutations in familial and sporadic amyotrophic lateral sclerosis Science 319, 1668-1672 10.1126/science.1154584

2. Lagier-Tourenne, C., Polymenidou, M., and Cleveland, D. W. (2010) TDP-43 and FUS/TLS: emerging roles in RNA processing and neurodegeneration Hum Mol Genet 19, 46,

3. Turner, M. R., Hardiman, O., Benatar, M., Brooks, B. R., Chio, A., de Carvalho, M. et al. (2013) Controversies and priorities in amyotrophic lateral sclerosis Lancet Neurol 12, 310–322 10.1016/S1474-4422(13)70036-X

4. Lee, E. B., Lee, V. M., and Trojanowski, J. Q. (2010) Gains or losses: molecular mechanisms of TDP43-mediated neurodegeneration Nat Rev Neurol 6, 211–220,

5. Xu, Z. S. (2012) Does a loss of TDP-43 function cause neurodegeneration? Mol Neurodegener 7, 27,

6. Buratti, E., and Baralle, F. E. (2008) Multiple roles of TDP-43 in gene expression, splicing regulation, and human disease Front Biosci 13, 867–878,

7. Yang, C., Tan, W., Whittle, C., Qiu, L., Cao, L., Akbarian, S., and Xu, Z. (2010) The C-terminal TDP-43 fragments have a high aggregation propensity and harm neurons by a dominant-negative mechanism PLoS One 5, e15878,

8. Kuo, P. H., Doudeva, L. G., Wang, Y. T., Shen, C. K., and Yuan, H. S. (2009) Structural insights into TDP-43 in nucleic-acid binding and domain interactions Nucleic Acids Res 37, 1799-1808,

9. Emanuele Buratti, F. E. B. (2001) Characterization and Functional Implications of the RNA Binding Properties of Nuclear Factor TDP-43, a Novel Splicing Regulator ofCFTR Exon 9 Journal of Biological Chemistry 276, 36337–36343,

10. Furukawa, Y., Suzuki, Y., Fukuoka, M., Nagasawa, K., Nakagome, K., Shimizu, H., et al. (2016) A molecular mechanism realizing sequence-specific recognition of nucleic acids by TDP-43 Sci Rep 6, 20576 10.1038/srep20576

11. Kuo, P. H., Chiang, C. H., Wang, Y. T., Doudeva, L. G., and Yuan, H. S. (2014) The crystal structure of TDP-43 RRM1-DNA complex reveals the specific recognition for UG- and TG-rich nucleic acids Nucleic Acids Res 42, 4712-4722 10.1093/nar/gkt1407

12. Mackness, B. C., Tran, M. T., McClain, S. P., Matthews, C. R., and Zitzewitz, J. A. (2014) Folding of the RNA Recognition Motif (RRM) Domains of the Amyotrophic Lateral Sclerosis (ALS)-linked Protein TDP-43 Reveals an Intermediate State J Biol Chem 289, 8264-8276,

13. Gershenson, A., Gierasch, L. M., Pastore, A., and Radford, S. E. (2014) Energy landscapes of functional proteins are inherently risky Nat Chem Biol 10, 884–891 10.1038/nchembio.1670

14. Korzhnev, D. M., Religa, T. L., and Kay, L. E. (2012) Transiently populated intermediate functions as a branching point of the FF domain folding pathway Proc Natl Acad Sci U S A 109, 17777–17782 10.1073/pnas.1201799109

15. Bilsel, M. A. E. K. C. K. C. R. M. M. I. O. (2007) Microsecond Hydrophobic Collapse in the Folding of Escherichia coli Dihydrofolate Reductase, an α/β-Type Protein journal of Molecular Biology 368, 219–229,

16. Rose, N. C. F. a. G. D. (2004) Reassessing random-coil statistics in unfolded proteins Proc Natl Acad Sci U S A 101, 12497–12502,

17. H. Jane Dyson, P. E. W. (2005) Elucidation of the Protein Folding Landscape by NMR Methods in Enzymology 394, 299–321,

18. Yulia Pustovalova, P. K., Michele Vendruscolo, and Dmitry M. Korzhnev (2015) Probing the Residual Structure of the Low Populated Denatured State of ADA2h under Folding Conditions by Relaxation Dispersion Nuclear Magnetic Resonance Spectroscopy Biochemistry 54, 4611–4622 doi.org/10.1021/acs.biochem.5b00345

19. Frédéric H.-T Allain, D. E. G., Philippe Bouvet, Juli Feigon (2000) Solution structure of the two N-terminal RNA-binding domains of nucleolin and NMR study of the interaction with its RNA target Journal of Molecular Biology 303, 227–241,

20. Marsh, J. A., Singh, V. K., Jia, Z., and Forman-Kay, J. D. (2006) Sensitivity of secondary structure propensities to sequence differences between alpha- and gamma-synuclein: implications for fibrillation Protein Sci 15, 2795-2804,

21. Qian Yi, M. L. S.-K., Eric J. Alm, David Baker (2000) NMR characterization of residual structure in the denatured state of protein L 299, 1341-1351,

22. Palmer, A. G., 3rd, Kroenke, C. D., and Loria, J. P. (2001) Nuclear magnetic resonance methods for quantifying microsecond-to-millisecond motions in biological macromolecules Methods Enzymol 339, 204-238,

23. Palmer, A. G. (2004) NMR characterization of the dynamics of biomacromolecules Chemical Reviews 104, 3623-3640, <Go to ISI>://000223290500008

24. Shen, Y., and Bax, A. (2010) SPARTAc in empirical NMR chemical shift prediction by means of an artificial neural network J Biomol NMR 48, 13–22,

25. Camilloni, C., De Simone, A., Vranken, W. F., and Vendruscolo, M. (2012) Determination of Secondary Structure Populations in Disordered States of Proteins Using Nuclear Magnetic Resonance Chemical Shifts Biochemistry 51, 2224-2231 10.1021/bi3001825

26. Kabsch W, S. C. (1983) Dictionary of protein secondary structure: pattern recognition of hydrogen-bonded and geometrical features Biopolymers 22, 2577-2637 10.1002/bip.360221211

27. Kathuria, S. V., Chan, Y. H., Nobrega, R. P., Özen, A., and Matthews, C. R. (2016) Clusters of isoleucine, leucine, and valine side chains define cores of stability in high-energy states of globular proteins: Sequence determinants of structure and stability Protein Science 25, 662–675,

28. Lu H, S. K. (1999) Steered molecular dynamics simulations of force-induced protein domain unfolding Proteins 35, 453–463,

29. Mu Gao, D. C., Viola Vogel, Klaus Schulten (2002) Identifying Unfolding Intermediates of FN-III10 by Steered Molecular Dynamics journal of Molecular Biology 323, 939–950 10.1016/S0022-2836(02)01001-X

30. Zhuravlev, P. I., Hinczewski, M., Chakrabarti, S., Marqusee, S., and Thirumalai, D. (2016) Force-dependent switch in protein unfolding pathways and transition-state movements Proc Natl Acad Sci U S A 113, E715–724 10.1073/pnas.1515730113

31. Tavella, D., Zitzewitz, J. A., and Massi, F. (2018) Characterization of TDP-43 RRM2 Partially Folded States and Their Significance to ALS Pathogenesis Biophys J 115, 1673-1680 10.1016/j.bpj.2018.09.011

32. Wang, Y.-T., Kuo, P.-H., Chiang, C.-H., Liang, J.-R., Chen, Y.-R., Wang, S. et al. (2013) The Truncated C-terminal RNA Recognition Motif of TDP-43 Protein Plays a Key Role in Forming Proteinaceous Aggregates J Biol Chem 288, 9049-9057,

33. Saini, A., and Chauhan, V. S. (2011) Delineation of the Core Aggregation Sequences of TDP-43 C-Terminal Fragment Chembiochem 12, 2495-2501 10.1002/cbic.201100427

34. Romano, M., Buratti, E., Romano, G., Klima, R., Del Bel Belluz, L., Stuani, C. et al. (2014) Evolutionarily conserved heterogeneous nuclear ribonucleoprotein (hnRNP) A/B proteins functionally interact with human and Drosophila TAR DNA-binding protein 43 (TDP-43) J Biol Chem 289, 7121-7130 10.1074/jbc.M114.548859

35. Ederle, H., Funk, C., Abou-Ajram, C., Hutten, S., Funk, E. B. E., Kehlenbach, R. H., et al. (2018) Nuclear egress of TDP-43 and FUS occurs independently of Exportin-1/CRM1 Sci Rep 8, 7084 10.1038/s41598-018-25007-5

36. Duan, L., Zaepfel, B. L., Aksenova, V., Dasso, M., Rothstein, J. D., Kalab, P., and Hayes, L. R. (2022) Nuclear RNA binding regulates TDP-43 nuclear localization and passive nuclear export Cell Rep 40, 111106 10.1016/j.celrep.2022.111106

37. Lukavsky, P. J., Daujotyte, D., Tollervey, J. R., Ule, J., Stuani, C., Buratti, E. et al. (2013) Molecular basis of UG-rich RNA recognition by the human splicing factor TDP-43 Nat Struct Mol Biol 20, 1443-1449 10.1038/nsmb.2698

38. Yan, X., Kuster, D., Mohanty, P., Nijssen, J., Pombo-Garcia, K., Rizuan, A., et al. (2024) Intra-condensate demixing of TDP-43 inside stress granules generates pathological aggregates bioRxiv 10.1101/2024.01.23.576837

39. Yu, H., Lu, S., Gasior, K., Singh, D., Vazquez-Sanchez, S., Tapia, O., et al. (2021) HSP70 chaperones RNA-free TDP-43 into anisotropic intranuclear liquid spherical shells Science 371, 10.1126/science.abb4309

40. Klaips, C. L., Jayaraj, G. G., and Hartl, F. U. (2018) Pathways of cellular proteostasis in aging and disease J Cell Biol 217, 51–63 10.1083/jcb.201709072

41. Igaz, L. M., Kwong, L. K., Xu, Y., Truax, A. C., Uryu, K., Neumann, M. et al. (2008) Enrichment of C-terminal fragments of TAR DNA-binding protein-43 cytoplasmic inclusions in brain but not in spinal cord of frontotemporal lobar degeneration and amyotrophic lateral sclerosis Am J Pathol 173, 182–194,

42. Uversky, V. N. (2015) Intrinsically disordered proteins and their (disordered) proteomes in neurodegenerative disorders Front Aging Neurosci 7, 18 10.3389/fnagi.2015.00018

43. Sgourakis, N. G., Yan, Y., McCullam, S., Wang, C., and Garcia, A. E. (2007) The Alzheimer’s peptides Aβ40 and 42 adopt distinct conformations in water: A combined MD / NMR study J Mol Biol 368, 1448-1457,

44. Benilova, I., Karran, E., and De Strooper, B. (2012) The toxic Abeta oligomer and Alzheimer’s disease: an emperor in need of clothes Nat Neurosci 15, 349–357 10.1038/nn.3028

45. Glabe, C. G., and Kayed, R. (2006) Common structure and toxic function of amyloid oligomers implies a common mechanism of pathogenesis Neurology 66, S74-S78,

46. Volles, M. J., and Landsbury, P. T. (2003) Zeroing in on the Pathogenic Form of α-Synuclein and Its Mechanism of Neurotoxicity in Parkinson’s Disease Biochemistry 42, 7871-7878 0.1021/bi030086j

47. Grad, L. I., Fernando, S. M., and Cashman, N. R. (2015) From molecule to molecule and cell to cell: prion-like mechanisms in amyotrophic lateral sclerosis Neurobiol Dis 77, 257–265 10.1016/j.nbd.2015.02.009

48. Jahn, T. R., and Radford, S. E. (2006) The Yin and Yang of protein folding The FEBS Journal 272, 5962-5970 10.1111/j.1742-4658.2005.05021.x

49. Karamanos, T. K., Pashley, C. L., Kalverda, A. P., Thompson, G. S., Mayzel, M., Orekhov, V. Y., and Radford, S. E. (2016) A Population Shift between Sparsely Populated Folding Intermediates Determines Amyloidogenicity J Am Chem Soc 138, 6271-6280 10.1021/jacs.6b02464

50. Khalil, B., Linsenmeier, M., Smith, C. L., Shorter, J., and Rossoll, W. (2024) Nuclear-import receptors as gatekeepers of pathological phase transitions in ALS/FTD Mol Neurodegener 19, 8 10.1186/s13024-023-00698-1

51. Guo, L., Mann, J. R., Mauna, J. C., Copley, K. E., Wang, H., Rubien, J. D. et al. (2023) Defining RNA oligonucleotides that reverse deleterious phase transitions of RNA-binding proteins with prion-like domains bioRxiv 10.1101/2023.09.04.555754

52. Pettersen, E. F., Goddard, T. D., Huang, C. C., Couch, G. S., Greenblatt, D. M., Meng, E. C., and Ferrin, T. E. (2004) UCSF Chimera--a visualization system for exploratory research and analysis J Comput Chem 25, 1605-1612,

53. Cavanagh, J., Fairbrother, W. J., Palmer, A. G., 3rd, and Skelton, N. J. (1996) Protein NMR Spectroscopy: Principles and Practice, Academic Press, San Diego

54. Henry, E., and Hofrichter, J. (1992) Singular value decomposition: Application to analysis of experimental data Methods Enzymol 210, 129–192,

55. Delaglio, F., Grzesiek, S., Vuister, G. W., Zhu, G., Pfeifer, J., and Bax, A. (1995) NMRPipe: a multidimensional spectral processing system based on UNIX pipes J Biomol NMR 6, 277–293,

56. Weiser, J., Senkin, P., and Still, W. C. (1999) Approximate atomic surfaces from linear combinations of pairwise overlaps (LCPO) J Comp Chem 20, 217–230,

57. Morgan, B. R., Zitzewitz, J. A., and Massi, F. (2017) Structural Rearrangement upon Fragmentation of the Stability Core of the ALS-Linked Protein TDP-43 Biophys J 113, 540–549 10.1016/j.bpj.2017.06.049

58. Phillips, J. C., Braun, R., Wang, W., Gumbart, J., Tajkhorshid, E., Villa, E. et al. (2005) Scalable molecular dynamics with NAMD Journal of Computational Chemistry 26, 1781-1802 10.1002/jcc.20289

59. Humphrey, W., Dalke, A., and Schulten, K. (1996) VMD – Visual Molecular Dynamics Journal of Molecular Graphics 14, 33–38,

60. Frishman, D., and Argos, P. (1995) Knowledge-based secondary structure assignment Proteins 23, 566–579,

61. Kathuria, S. V., Day, I. J., Wallace, L. A., and Matthews, C. R. (2008) Kinetic Traps in the folding of βα-Repeat Proteins: CheY Initially Misfolds before Accessing the Native Conformation J Mol Biol 382, 467–484,

62. Wu, Y., Vadrevu, R., Kathuria, S., Yang, X., and Matthews, C. R. (2007) A tightly packed hydrophobic cluster directs the formation of an off-pathway sub-millisecond folding intermediate in the α subunit of tryptophan synthase, a TIM barrel protein J Mol Biol 366, 1624-1638,

63. Fernandez-Escamilla, A. M., Rousseau, F., Schymkowitz, J., and Serrano, L. (2004) Prediction of sequence-dependent and mutational effects on the aggregation of peptides and proteins Nat Biotechnol 22, 1302-1306 10.1038/nbt1012

64. Romero P, O. Z., Li X, Garner EC, Brown CJ, Dunker AK (2001) Sequence complexity of disordered protein Proteins 42, 38–48 10.1002/1097-0134(20010101)42:1<38::aid-prot50>3.0.co;2-

