## Supplemental Material for "A hydrophobic core stabilizes the residual structure in the RRM2 intermediate state of the ALS-linked protein TDP-43"

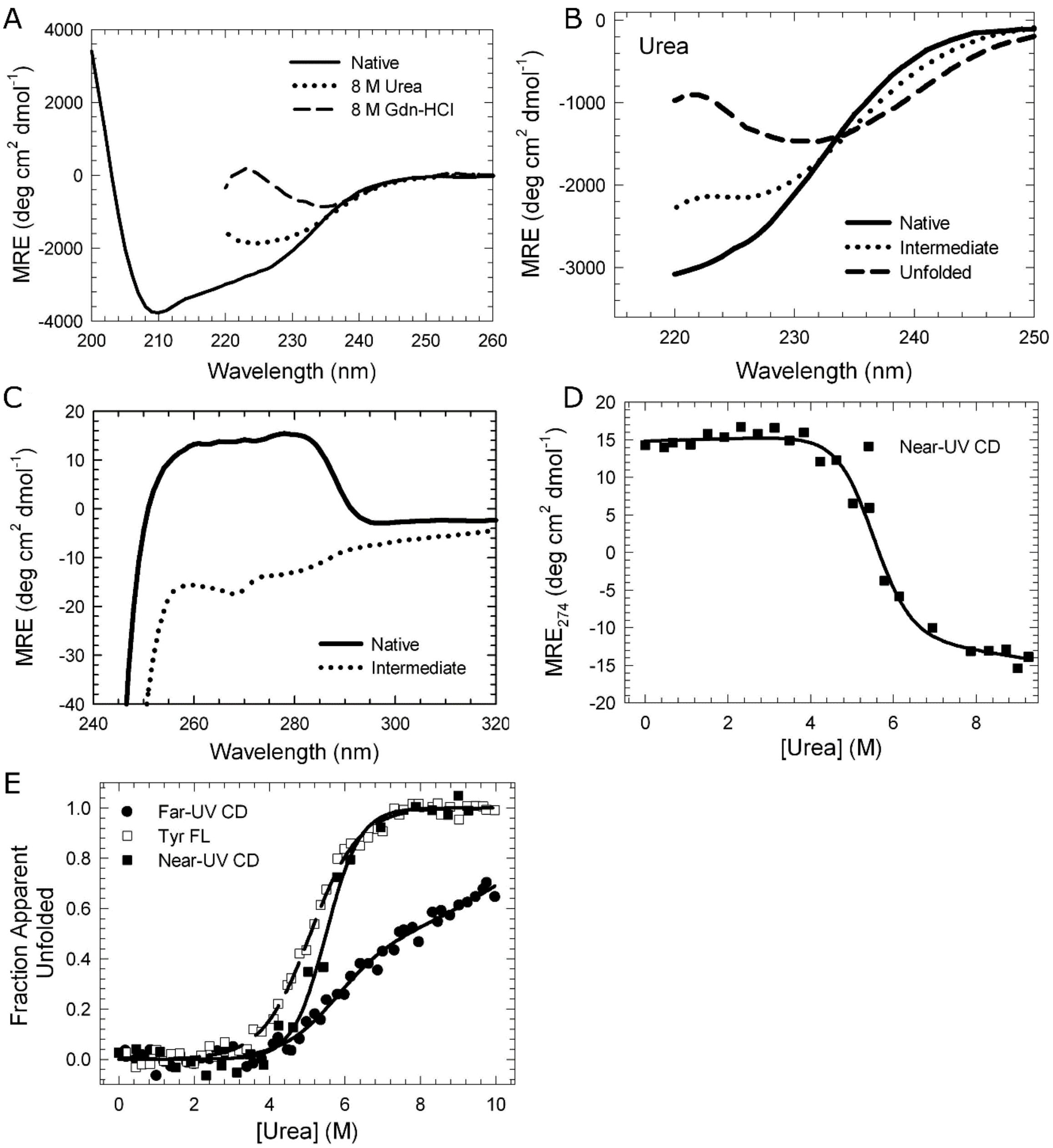


**Figure S1. Chemical denaturation studies of RRM2 reveal the presence of an intermediate in the unfolding pathway.** (A) Far-UV CD spectra of RRM2 under native conditions (solid) and denaturing conditions, incubated with either 8 M urea (dotted) or 8 M Gdn-HCl (dashed). (B) SVD of the far-UV CD wavelength scans as a function of urea was used to obtain spectra of the native (solid), intermediate (dotted) and unfolded (dashed) component species. (C) Near-UV CD spectrum of the native (solid) and intermediate (dotted) states of RRM2. (D) Global modeling of the near-UV spectra as a function of urea to a two-state mechanism (N⇆I/U). The mean residue ellipticity at 274 nm is shown (squares) with the global fit (solid line) of all wavelengths from 250-285 nm. (E) Fraction apparent unfolded profiles by CD (closed circles), Tyr FL (open circles) and near-UV CD (open diamonds). Fits of the data are shown with solid or dashed lines.

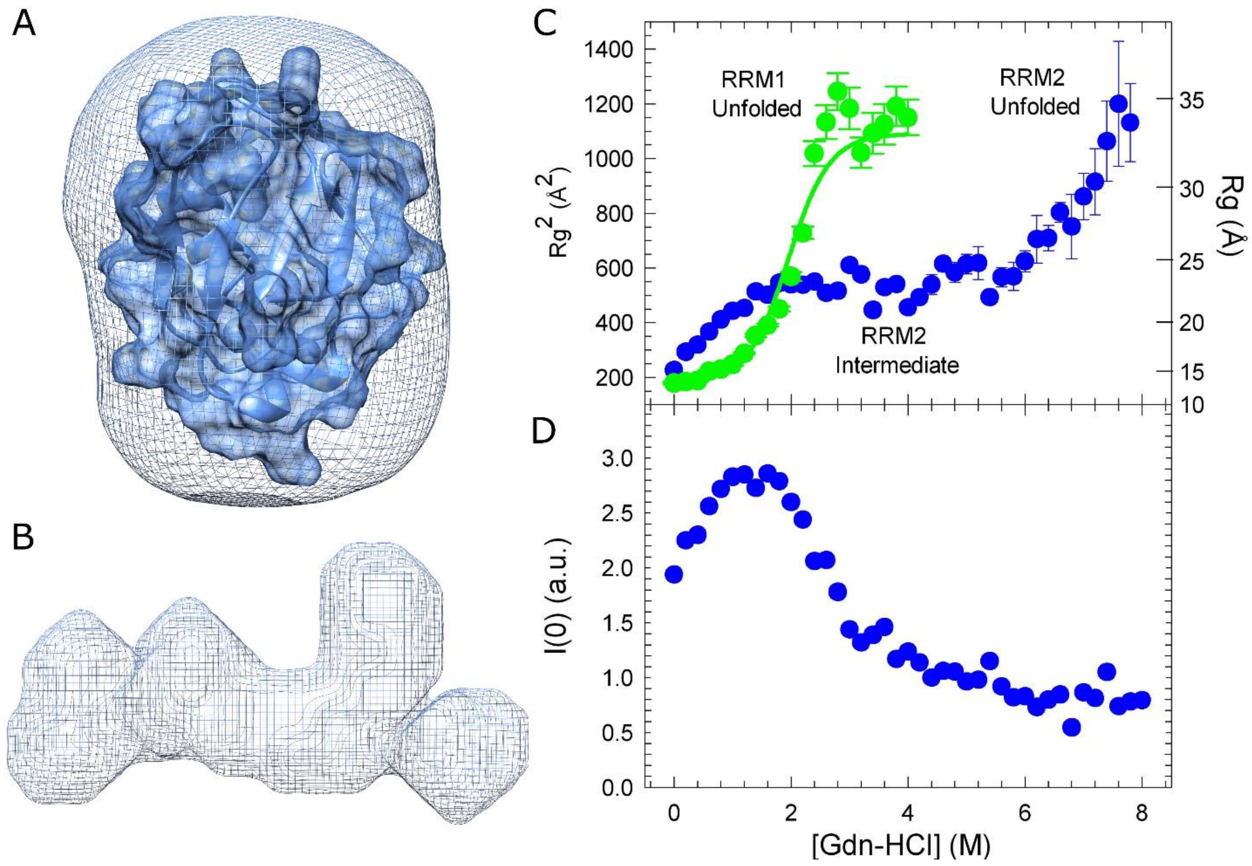


**Figure S2. RRM2 loses the majority of its globular structure in the intermediate state.** (A) The electron density envelope of the native state of RRM2 obtained by LC-SAXS overlaid with the NMR structure of RRM2 (pdb: 1wf0). (B) The electron density envelope of a representative RRM2 intermediate state at 8 M urea. (C) Equilibrium unfolding profile of RRM1 (green) and RRM2 (blue) monitored by SAXS using Gdn-HCl as a denaturant. RRM1 undergoes a two-state transition upon addition of Gdn-HCl to populate the RRM1 unfolded state. At low concentrations of Gdn-HCl, RRM2 expands from 15 Å to 23 Å (Table S2) and maintains this size conformation until 6 M Gdn-HCl. At concentrations > 6 M Gdn-HCl, the unfolded state dominates the scattering profile with an Rg similar to the theoretical value of 28 Å. (D) The plot of the intensity at zero scattering angle, I(0), at each urea concentration for RRM2 (blue). I(0) is non-linear for RRM2 with increased I(0) at < 2 M Gdn-HCl, suggesting the expanded conformation at these concentrations are a result of sample aggregation and not the population of the RRM2 intermediate.

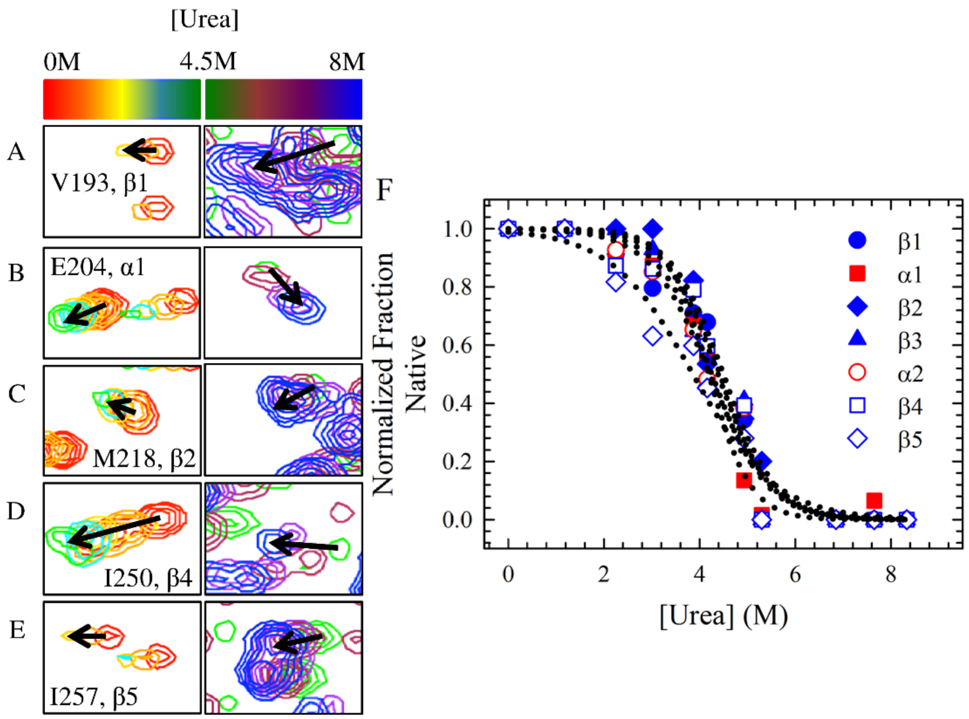


**Fig. S3. Characterization of the residual structure in the RRM2 intermediate state.** (A-E) HSQC cross-peaks of one residue from each secondary structural element in the native (left) and intermediate (right) baselines of the RRM2 equilibrium unfolding profile monitored by NMR. (A) V193: β1, (B) E204: α1, (C) M218: β2, (D) I250: β4, and (E) I257: β5. Chemical shift perturbations start from the native state (red) and conclude at the intermediate state (blue), where the black arrow indicates the direction of the change. (F) Equilibrium unfolding profiles for each secondary structural element, β1 (closed circles), α1 (closed squares), β2 (closed diamonds), β3 (closed triangles), α2 (open circles), β4 (open squares) and β5 (open diamonds), which unfold by a two-state mechanism (N⇆I/U).

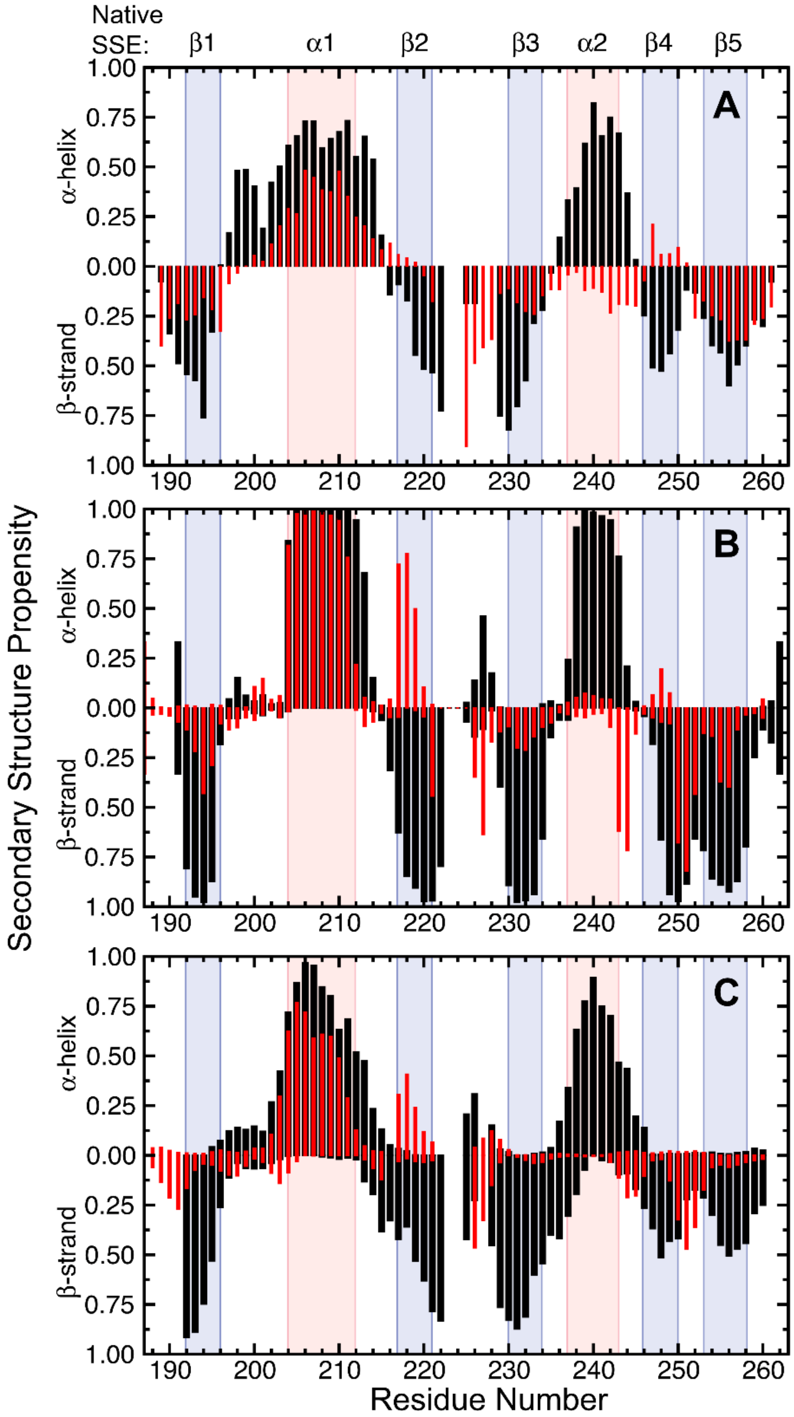


**Figure S4. RRM2 secondary structure propensity in the intermediate state is different from the native state.** Secondary structure propensity of the RRM2 native (black bars) and intermediate (red bars) states obtained from the $^{1}H$, $^{15}N$, Cα and Cβ chemical shifts deviations from random coil. Secondary structure propensity was calculated using three different algorithms: (A) SSP (Marsh et al., 2006), (B) SPARTA+ (Shen and Bax, 2010) and (C) δ2D (Camilloni et al., 2012). Positive and negative values represent α-helical and β-strand propensity, respectively. Pink (α-helices) and light blue (β-strand) boxes highlight the secondary structural elements observed in the native structure.

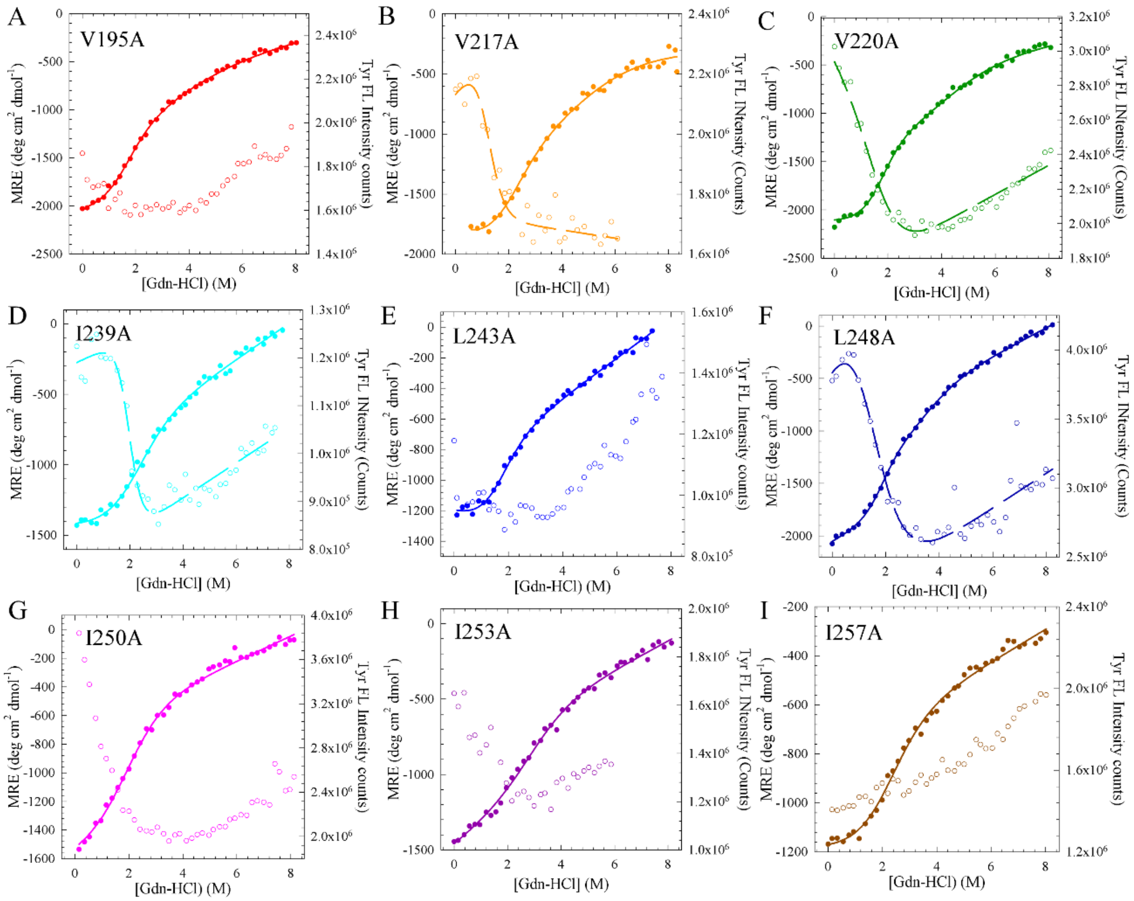


**Figure S5. Mutational analysis of the RRM2 shows destabilization of the hydrophobic core.** Equilibrium unfolding profiles of the RRM2 ILV Mutants by CD (closed circles) and Tyr FL (open circles).

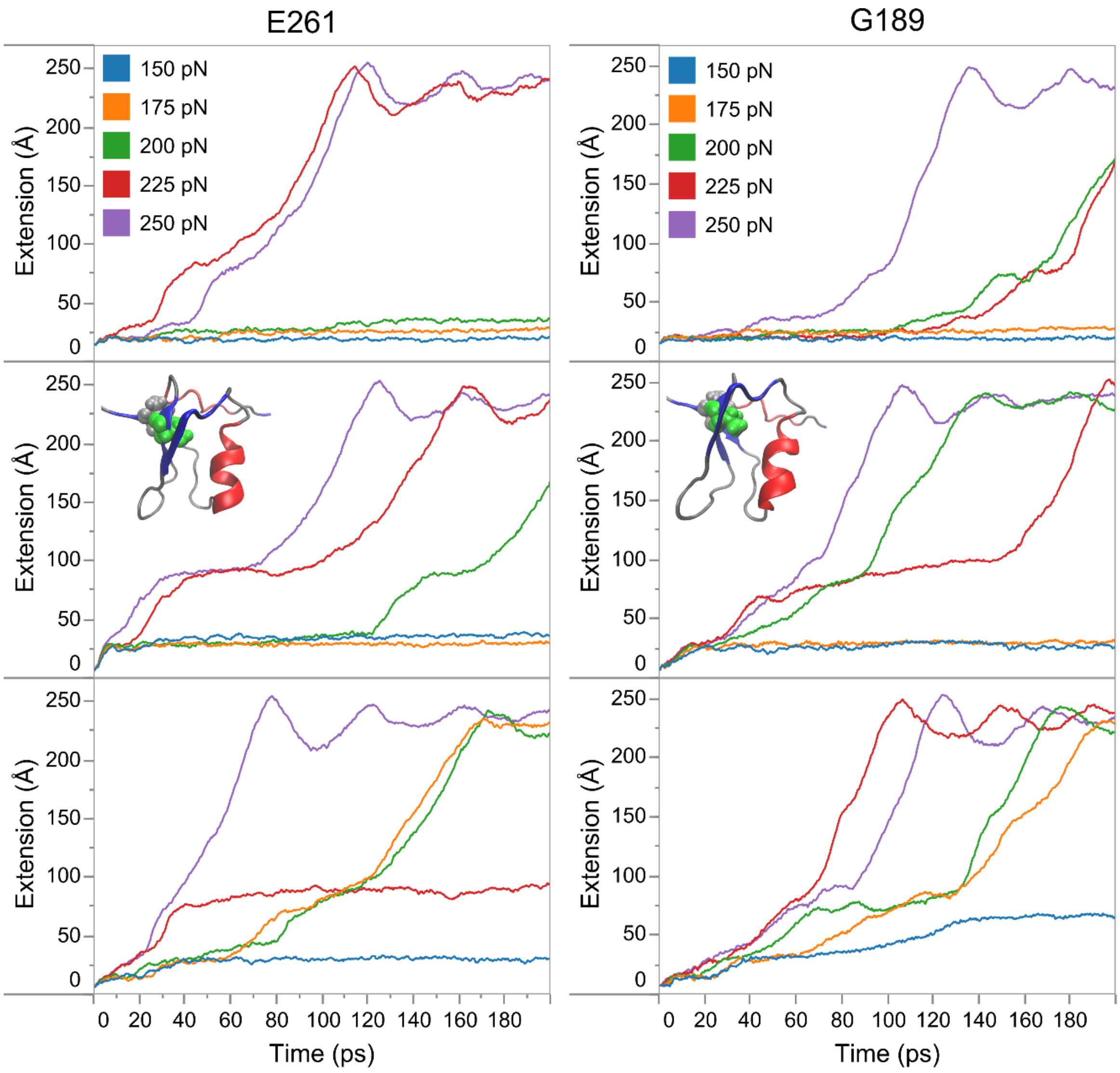


**Figure S6. Extension profiles are consistent across different initial conditions and pulling atom.** Extension profiles as a function of time for constant forces ranging from 150 to 250 pN. Profiles are labeled by the atom being pulled (E261 or G189) and start from three different initial conditions (top, middle, bottom). E261, middle corresponds to the initial conditions shown in the main paper in Fig. 6. A representative structure from the plateau region of the second initial condition (middle) pulled at E261 or G189 with a constant force of 225 pN (red) is shown. Gray ribbon shows RRM2 backbone of the remaining structured elements with β strands in blue and α helices in red. Van der Waals representations are shown of V193 (gray) and V232 (green).

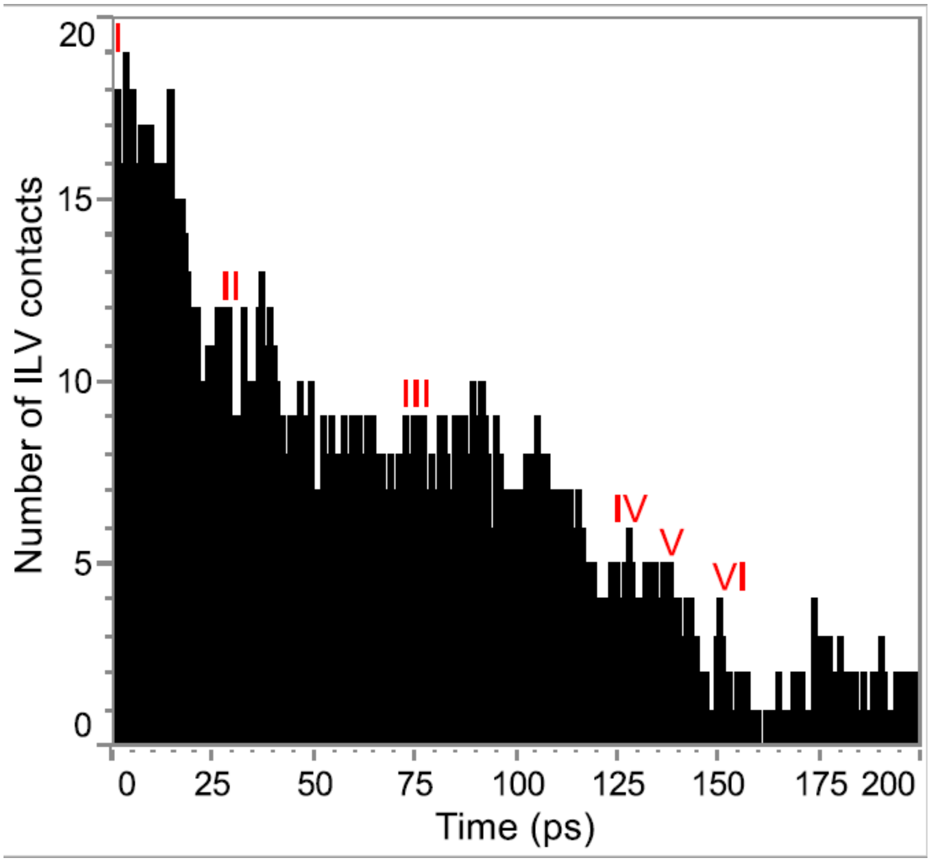


**Figure S7. Total number of ILV contacts decreases along the SMD unfolding pathway.** Number of contacts between ILV residues as a function of time for the unfolding pathway under 225 pN of constant force. Roman numerals (I-VI) mark the locations of the representative structures shown in Fig. 6, where III marks the location of the proposed intermediate structure.

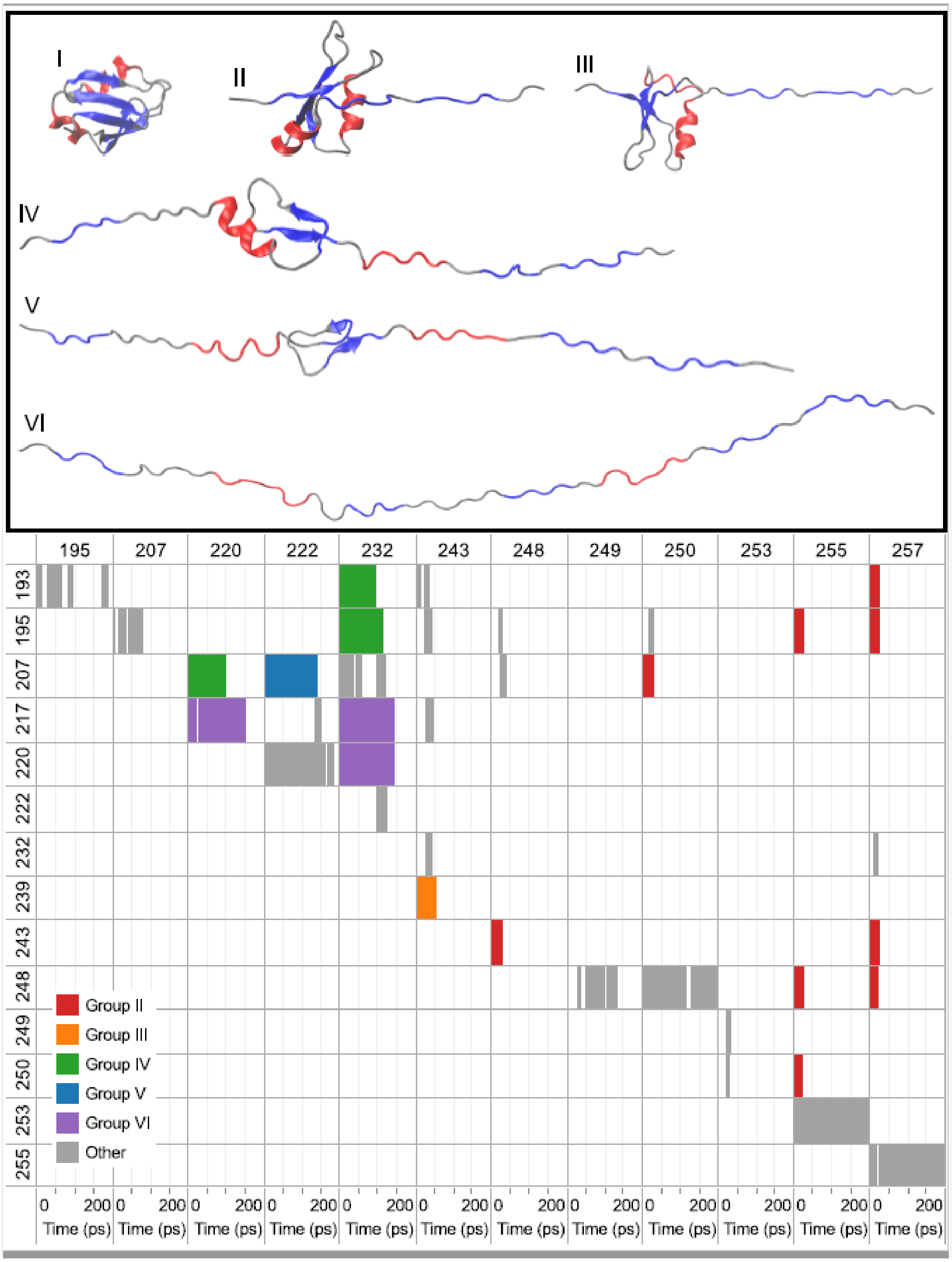


**Figure S8. Timeline of the SMD unfolding pathway highlights critical contacts in each conformational state.** (Top) Representative structures (I-VI) of the unfolding pathway under 225 pN of force. Gray ribbon shows RRM2 backbone with β-strands in blue and α-helices in red. (Bottom) Hydrophobic contacts between ILV residues as a function of time in the unfolding pathway under 225 pN of constant force. Colored bars indicate that a hydrophobic contact is present between a pair of residues. Groups II-VI correspond to groups of contacts that have been lost in the sample conformations II-VI in the top panel. Transient contacts or contacts between neighboring residues that are not lost prior to the fully extended conformation (VI) are colored gray.

| A | $R_{gyration}$ (Å) | SASA (Å$^2$) |
|---|---|---|
| NMR structure | 11.3 | 4803 |
| Implicit solvent | 11.5 ± 0.1 | 4784 ± 101 |
| Explicit solvent | 11.4 ± 0.1 | 4750 ± 114 |

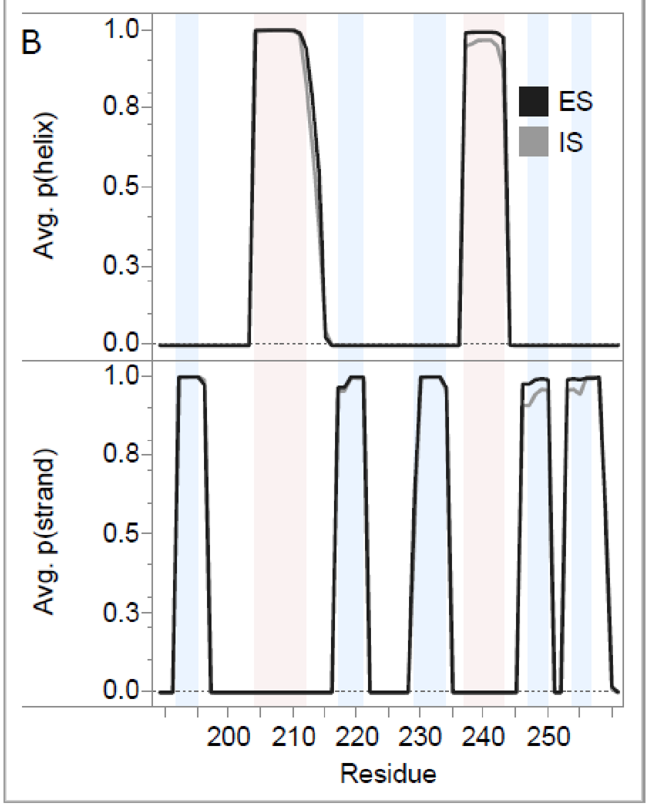


**Figure S9. RRM2 MD simulations in explicit and implicit solvent capture the same secondary structure.** (A) Comparison of structural metrics between solvent models. Mean and standard deviation of the radius of gyration ($R_{gyration}$) and solvent accessible surface area (SASA) for RRM2 of the NMR structure (pdb: 1wf0) and from implicit and explicit solvent simulations. (B) Comparison of secondary structural probabilities from simulations in explicit solvent (ES: black) and implicit solvent (IS: gray). Shading indicates the location of α-helices (pink) and β-strands (light blue) in the NMR structure (pdb: 1wf0). The dotted gray zero line is shown for visual clarity. The secondary structural probabilities were calculated using the combined average of three trajectories for a total production time of 540 ns (ES) and 720 ns (IS).

### Supplemental Tables

| | Rg (Å) | | |
|---|---|---|---|
| Denaturant | Native | Intermediate | Unfolded |
| Urea | 13.4 ± 0.3 | 23.4 ± 0.4 | *N/D |
| Gdn-HCl | 15.0 ± 0.1 | 23.1 ± 0.3 | 28.5 ± 1.2 |

**Table S1. Comparison of the dimensions of the RRM2 native, intermediate and unfolded states in urea and Gdn-HCl**. *N/D indicates the Rg of the unfolded state was not determined in urea because the unfolded state is not significantly populated at high urea concentrations.

| Secondary Structure Element | ΔG° | m | $C_m$ |
|---|---|---|---|
| β1 | 4.64 ± 0.89 | 1.06 ± 0.20 | 4.38 ± 1.19 |
| α1 | 5.61 ± 0.73 | 1.34 ± 0.17 | 4.19 ± 0.76 |
| β2 | 4.80 ± 0.58 | 1.07 ± 0.13 | 4.50 ± 0.77 |
| β3 | 4.29 ± 0.83 | 1.01 ± 0.19 | 4.25 ± 1.16 |
| α2 | 3.88 ± 0.67 | 0.92 ± 0.16 | 4.21 ± 1.02 |
| β4 | 4.95 ± 0.97 | 1.12 ± 0.22 | 4.42 ± 1.22 |
| β5 | 2.61 ± 0.45 | 0.68 ± 0.11 | 3.86 ± 0.93 |

**Table S2. NMR thermodynamic parameters for each secondary structural element in RRM2.** From the urea titrations, the normalized fraction native of each secondary structure was modeled to a two-state mechanism (N⇆I/U) to obtain the free energy of unfolding (ΔG°), m-value, and midpoint ($C_m$) for the transition. Units: ΔG° (kcal $mol^{-1}$), m (kcal $mol^{-1}$ $M^{-1}$) and $C_m$ (M). The β-strands and α-helices in the native state of RRM2 are colored blue and red, respectively.

| | N ⇄ U | | |
|---|---|---|---|
| **Variant** | **ΔG°** | **m** | **$C_m$** |
| I239A | 1.90 ± 0.06 | 0.89 ± 0.03 | 2.13 ± 0.10 |
| L243A | 1.94 ± 0.03 | 1.22 ± 0.05 | 1.59 ± 0.07 |
| L248A | 2.22 ± 0.04 | 1.10 ± 0.06 | 2.02 ± 0.12 |
| I250A | 1.64 ± 0.02 | 0.84 ± 0.01 | 1.93 ± 0.03 |
| I253A | 2.22 ± 0.05 | 0.75 ± 0.06 | 2.96 ± 0.25 |
| I257A | 1.88 ± 0.04 | 0.87 ± 0.03 | 2.16 ± 0.10 |

**Table S3. Thermodynamic parameters of the Class I mutations modeled to a two-state equilibrium unfolding profile (N ⇄ U).** ΔG: free energy (kcal $mol^{-1}$), m: buried surface area (kcal $mol^{-1}$ $M^{-1}$), and $C_m$: midpoint (M).

| | N ⇄ I | | | I ⇄ U | | | | | |
|---|---|---|---|---|---|---|---|---|---|
| **Variant** | **ΔG°$_1$** | **$m_1$** | **$C_{m1}$** | **ΔG°$_2$** | **$m_2$** | **$C_{m2}$** | **ΔG°$_{total}$** | **$m_{total}$** | **I (%)** |
| WT | 3.60 ± 0.15 | 1.30 ± 0.06 | 2.77 ± 0.17 | 3.82 ± 0.23 | 0.71 ± 0.03 | 5.38 ± 0.40 | 7.42 ± 0.27 | 2.01 ± 0.07 | 0.2 |
| V195A | 1.79 ± 0.03 | 1.03 ± 0.01 | 1.74 ± 0.03 | 1.35 ± 0.05 | 0.41 ± 0.03 | 3.29 ± 0.29 | 3.14 ± 0.06 | 1.44 ± 0.03 | 4.4 |
| V217A | 1.75 ± 0.03 | 0.84 ± 0.08 | 2.08 ± 0.20 | 1.82 ± 0.03 | 0.44 ± 0.04 | 4.13 ± 0.38 | 3.57 ± 0.04 | 1.28 ± 0.09 | 4.7 |
| V220 | 2.12 ± 0.04 | 1.12 ± 0.02 | 1.89 ± 0.04 | 1.91 ± 0.03 | 0.50 ± 0.02 | 3.82 ± 0.16 | 4.03 ± 0.05 | 1.62 ± 0.03 | 2.6 |

**Table S4. Thermodynamic parameters of the Class II mutations modeled to a three-state equilibrium unfolding profile (N ⇄ I ⇄ U).** ΔG: free energy (kcal mol$^{-1}$), m: buried surface area (kcal mol$^{-1}$ M$^{-1}$), $C_m$: midpoint (M), and I: population of intermediate state under native conditions at pH 7.2 and 20°C determined from thermodynamic parameters (percentage).
